# The impact of CO_2_/HCO_3_^-^ availability on anaplerotic flux in PDHC-deficient *Corynebacterium glutamicum* strains

**DOI:** 10.1101/663856

**Authors:** Aileen Krüger, Johanna Wiechert, Cornelia Gätgens, Tino Polen, Regina Mahr, Julia Frunzke

**Affiliations:** Institut für Bio- und Geowissenschaften, IBG-1: Biotechnology, Forschungszentrum Jülich, 52425 Jülich, Germany; SenseUp GmbH, c/o Campus Forschungszentrum Jülich, 52425 Jülich, Germany

**Author notes:** A.K. and J.W. contributed equally to this work. Address correspondence to Julia Frunzke, and Regina Mahr,.

## Abstract

The pyruvate dehydrogenase complex (PDHC) catalyzes the oxidative decarboxylation of pyruvate yielding acetyl-CoA and CO_2_. The PDHC-deficient *Corynebacterium glutamicum* strain Δ*aceE* is therefore lacking an important decarboxylation step in central metabolism. Additional inactivation of *pyc*, encoding pyruvate carboxylase, resulted in a >15 hour lag phase in the presence of glucose, while no growth defect was observed on gluconeogenetic substrates like acetate. Growth was successfully restored by deletion of *ptsG* encoding the glucose-specific permease of the PTS system, thereby linking the observed phenotype to the increased sensitivity of strain Δ*aceE* Δ*pyc* to glucose catabolism. In the following, strain Δ*aceE* Δ*pyc* was used to systematically study the impact of perturbations of the intracellular CO_2_/HCO_3_^-^ pool on growth and anaplerotic flux. Remarkably, all measures leading to enhanced CO_2_/HCO_3_^-^ levels, such as external addition of HCO_3_^-^, increasing the pH, or rerouting metabolic flux via pentose phosphate pathway, at least partially eliminated the lag phase of strain Δ*aceE* Δ*pyc* on glucose medium. In accordance, inactivation of the urease enzyme, lowering the intracellular CO_2_/HCO_3_^-^ pool, led to an even longer lag phase accompanied with the excretion of L-valine and L-alanine. Transcriptome analysis as well as an adaptive laboratory evolution experiment of strain Δ*aceE* Δ*pyc* revealed the reduction of glucose uptake as a key adaptive measure to enhance growth on glucose/acetate mixtures. Altogether, our results highlight the significant impact of the intracellular CO_2_/HCO_3_^-^ pool on metabolic flux distribution, which becomes especially evident in engineered strains suffering from low endogenous CO_2_ production rates as exemplified by PDHC-deficient strains.

**Importance:** CO_2_ is a ubiquitous product of cellular metabolism and an essential substrate for carboxylation reactions. The pyruvate dehydrogenase complex (PDHC) catalyzes a central metabolic reaction contributing to the intracellular CO_2_/HCO_3_^-^ pool in many organisms. In this study, we used a PDHC-deficient strain of *Corynebacterium glutamicum*, which was additionally lacking pyruvate carboxylase (Δ*aceE* Δ*pyc*). This strain featured a >15 h lag phase during growth on glucose-acetate mixtures. We used this strain to systematically assess the impact of alterations in the intracellular CO_2_/HCO_3_^-^ pool on growth on glucose-containing medium. Remarkably, all measures enhancing the CO_2_/HCO_3_^-^ levels successfully restored growth emphasizing the strong impact of the intracellular CO_2_/HCO_3_^-^ pool on metabolic flux especially in strains suffering from low endogenous CO_2_ production rates.

## Introduction

CO_2_ is an inevitable product and at the same time an essential substrate of microbial metabolism. In water, CO_2_ is in equilibrium with HCO_3_^-^ and CO_3_^2-^, which is influenced by the pH of the medium (1). As a product of decarboxylating reactions and substrate for carboxylations, CO_2_ and bicarbonate (HCO_3_^-^) are involved in various metabolic processes. Consequently, the intracellular CO_2_/HCO_3_^-^ pool has a significant impact on metabolic fluxes like e.g. anaplerosis, especially in the early phase of cultivations when cell density is low (2).

During aerobic growth, the pyruvate dehydrogenase complex (PDHC), which is conserved in various microbial species, catalyzes an important reaction contributing to the intracellular CO_2_/HCO_3_^-^ pool (3, 4). The PDHC is a multienzyme complex belonging to the family of 2-oxo acid dehydrogenase complexes also comprising the 2-ketoglutarate dehydrogenase complex (ODHC) and the branched-chain 2-ketoacid dehydrogenase (3, 5). In particular, PDHC catalyzes the oxidative decarboxylation of pyruvate to acetyl-CoA, thereby producing one molecule CO_2_ and one molecule NADH per mol pyruvate.

The Gram-positive actinobacterium *Corynebacterium glutamicum* represents an important platform strain used in industrial biotechnology for the production of amino acids, proteins and various other value-added products (6–10). In this organism, the PDHC complex represents a central target to engineer metabolic pathways of pyruvate-derived products including L-valine, isobutanol, ketoisovalerate and L-lysine. To this end, different studies focused on the reduction or complete abolishment of PDHC activity to improve precursor availability (4, 11–13). Due to the deficiency of PDHC activity, however, cells are not able to grow on glucose as single carbon source which can be circumvented by the addition of acetyl-CoA refuelling substrates such as acetate. In presence of both carbon sources glucose and acetate, PDHC-deficient strains initially form biomass from acetate and subsequently, convert glucose into products (e.g. L-valine or L-alanine) in the stationary phase (12). In contrast to *Escherichia coli* or *Bacillus subtilis*, *C. glutamicum* does not normally show the typical diauxic growth behaviour, but prefers co-metabolization of many carbon sources (10, 14). Co-utilization of glucose and acetate has been studied in detail, showing that consumption rates of both carbon sources decrease, but the total carbon consumption is comparable to growth on either carbon source alone which is regulated by SugR activity (15). Further examples for co-metabolization of glucose with other sugars or organic acids include fructose (16), lactate (17) as well as pyruvate (14) and gluconate (18). It was shown, that upon growth on glucose-acetate mixtures, the glyoxylate shunt is required to fuel the oxaloacetate pool in the wild type (15). Here it is important to note that the glyoxylate shunt is active in the presence of acetate, but repressed by glucose (19, 20).

For growth on glycolytic carbon sources, organisms depend on TCA cycle replenishing reactions constituting the anaplerotic node (or phosphoenolpyruvate-pyruvate-oxaloacetate node), comprised of different carboxylating and decarboxylating reactions (21). Consequently, flux via these reactions is influenced by the intracellular CO_2_/HCO_3_^-^ pool. In contrast to most other organisms, *C. glutamicum* possesses both anaplerotic carboxylases namely the phosphoenolpyruvate carboxylase (PEPCx) (EC 4.1.1.31) encoded by the *ppc* gene (22, 23) and the pyruvate carboxylase (PCx) (EC 6.4.1.1) encoded by *pyc* (21, 24). These two C3-carboxylating enzymes catalyze bicarbonate-dependent reactions yielding oxaloacetate from PEP or pyruvate, respectively. In *C. glutamicum*, it was shown that these two enzymes could replace each other to a certain extent depending on the intracellular concentrations of the respective effectors for each enzyme. However, under standard conditions during growth on glucose, the biotin-containing PCx contributes to the main anaplerotic activity of 90% compared to PEPCx (24). Remarkably, the Michaelis-Menten constants of both carboxylases are about 30-fold higher in comparison to *Escherichia coli* PEP carboxylase (25). Apparently, a low CO_2_/HCO_3_^-^ pool may thus limit anaplerotic flux, which is of special relevance in early phases when biomass concentration is low, but high aeration may strip dissolved CO_2_ from the medium.

Several studies revealed the inhibitory effect of low p_CO2_ on microbial growth (2, 26, 27). In the case of *C. glutamicum*, low p_CO2_ pressures led to a significant drop in growth rate in turbidostatic continuous cultures (28) as well as batch cultures (29). It was further demonstrated that the reduced flux through anaplerotic reactions under low CO_2_/HCO_3_^-^ levels led to an increased production of the pyruvate-derived amino acids L-alanine and L-valine (29).

While several previous studies focused on the impact of CO_2_ by altering the CO_2_ partial pressure in the process, we applied targeted genetic perturbations to systematically assess the impact of the intracellular CO_2_/HCO_3_^-^ pool on anaplerotic flux and growth of *C. glutamicum*. Here, we focused on PDHC-deficient strains suffering from a reduced intracellular CO_2_/HCO_3_^-^ pool during growth on glucose-acetate mixtures. The effect of low CO_2_/HCO_3_^-^ levels became even more evident, upon additional deletion of the *pyc* gene encoding the dominant anaplerotic enzyme leading to a drastically elongated lag phase of a *C. glutamicum* Δ*aceE* Δ*pyc* strain during growth on glucose acetate mixtures. This effect could be attributed to a reduced tolerance of these strains towards glucose and was successfully complemented by deletion of *ptsG* encoding the glucose-specific permease of the PTS system. Remarkably, growth was successfully restored by increasing the intracellular CO_2_/HCO_3_^-^ pool by the addition of bicarbonate, by increasing the pH, by re-routing metabolic flux over the pentose phosphate pathway or by refueling of the oxaloacetate pool by the addition of TCA intermediates to the growth medium. Finally, an adaptive laboratory evolution (ALE) of *C. glutamicum* Δ*aceE* Δ*pyc* strain on minimal medium containing glucose and acetate revealed the abolishment of glucose uptake as a primary strategy to allow for growth on acetate.

## Results

### A lowered intracellular CO_2_/HCO_3_^-^ pool impacts growth of PDHC-deficient strains

The pyruvate dehydrogenase complex (PDHC) comprises a key metabolic reaction in the central metabolism by catalyzing the oxidative decarboxylation of pyruvate to acetyl-CoA (Fig. 1), thereby producing one molecule CO_2_ and one molecule NADH per mol pyruvate. Previous studies already revealed that PDHC-deficient strains feature an about two-fold decreased excretion of CO_2_ during exponential growth (30). It is therefore likely to assume that a reduction of the intracellular CO_2_/HCO_3_^-^ pool has an impact on metabolic flux distribution in PDHC-deficient strains. A simple growth comparison of the C*. glutamicum* wild type and a strain lacking the Δ*aceE* gene (thus PDHC-deficient) revealed a slightly increased lag phase of the Δ*aceE* strain which especially became evident when cells were grown in microtiter plates, while the maximal growth rate appeared to be unaffected (Δ*aceE*: 0.40 ± 0.01 h^-1^; WT: 0.41 ± 0.01 h^-1^) (Fig. 2A). Remarkably, this lag phase was completely eliminated by the addition of 100 mM HCO_3_^-^ attributing the effect to the lowered intracellular CO_2_/HCO_3_^-^ pool of this mutant (Fig. 2B).

**Figure 1:**
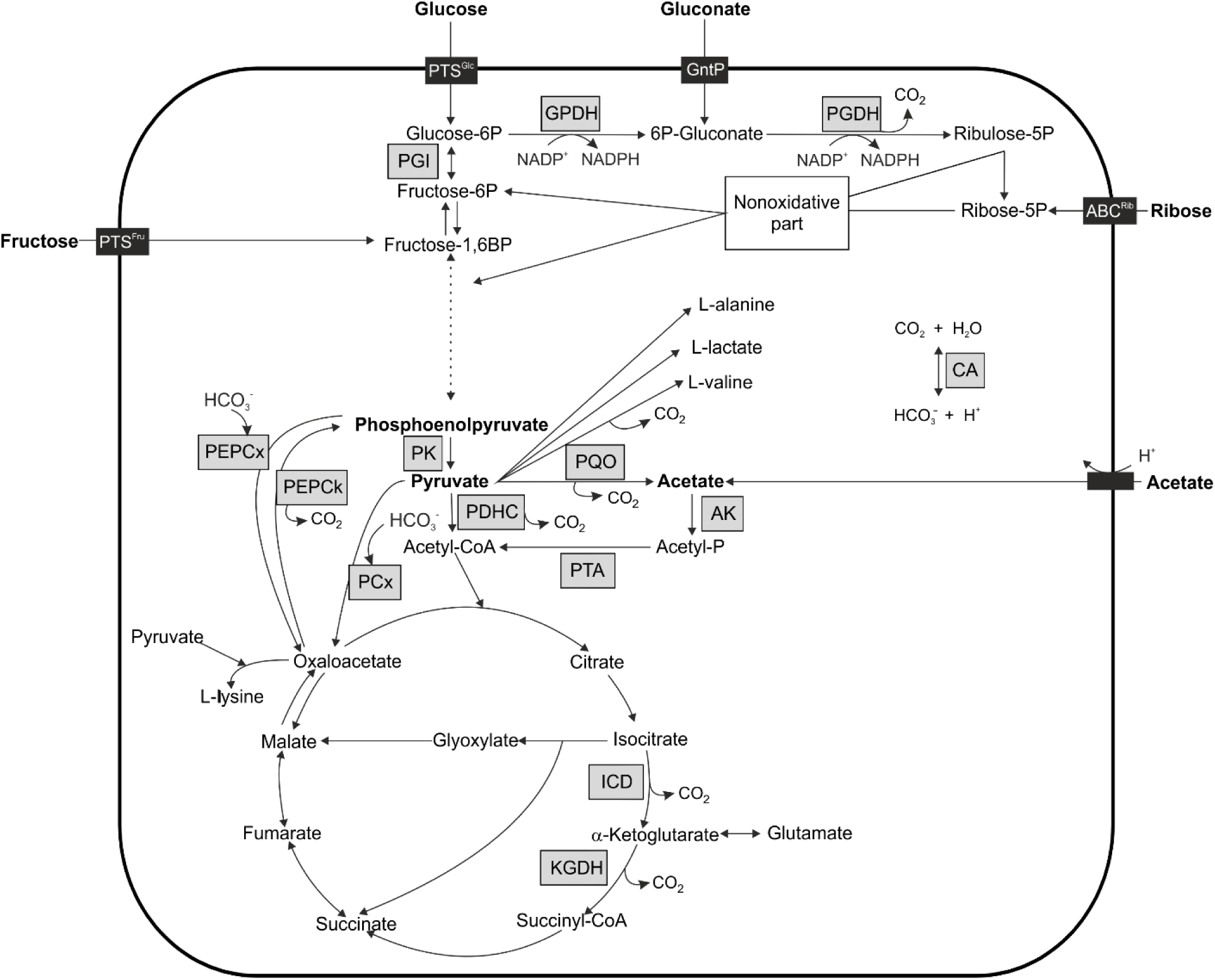
Schematic overview of the central carbon metabolism of *C. glutamicum.* The main aspects of glycolysis, gluconeogenesis, pentose phosphate pathway, tricarboxylic acid cycle (TCA) and anaplerosis are shown, while grey boxes represent relevant enzymes. Carboxylation as well as decarboxylation steps are given. ABC^Rib^ = ATP-binding cassette transporter for ribose, AK = acetate kinase, CA = carbonic anhydrase, GntP = gluconate permease, GPDH = glucose-6P dehydrogenase, ICD = Isocitrate dehydrogenase, KGDH = α-ketoglutarate dehydrogenase complex, PCx = pyruvate carboxylase, PDHC = pyruvate dehydrogenase complex, PEPCk = PEP carboxykinase, PEPCx = PEP carboxylase, PGDH = 6-P gluconate dehydrogenase, PGI = phosphoglucose isomerase, PK = pyruvate kinase, PQO = pyruvate:quinone oxidoreductase, PTA = phosphotransacetylase, PTS^Glc^ = permease of phosphotransferase system for glucose, PTS^Fru^= permease of phosphotransferase system for fructose.

**Figure 2:**
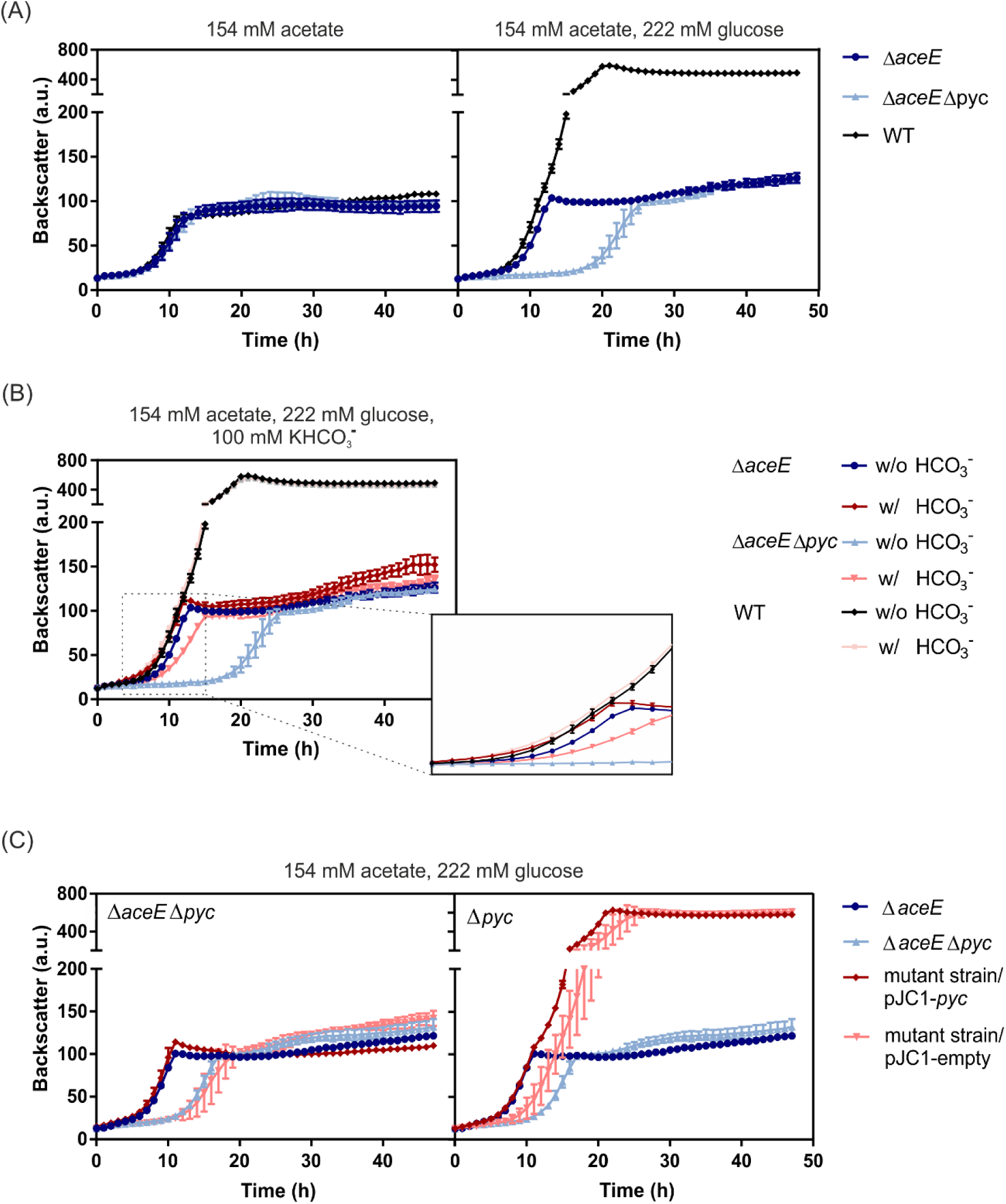
Impact of glucose on *aceE* and *pyc* mutant strains. Growth curves shown are based on the backscatter measurements in a microtiter cultivation system, inoculated at an OD_600_ of 1. Data represent the average of three biological replicates, error bars the standard deviation. (A) The *C. glutamicum* strains Δ*aceE* and Δ*aceE* Δ*pyc*, as well as the wild type were inoculated in CGXII media containing 154 mM acetate (left) or 154 mM acetate plus 222 mM glucose (right). (B) Strains Δ*aceE* and Δ*aceE* Δ*pyc* as well as WT cells were cultivated in CGXII medium containing 154 mM acetate and 222 mM glucose and supplemented either with (red lines) or without 100 mM KHCO_3_^-^ (blue and black lines). The box zooms in the time interval between 4 h and 15 h. (C) Strains Δ*aceE* Δ*pyc* (left) and Δ*pyc* (right) were transformed with pJC1-*pyc* plasmid for complementation of the *pyc* deletion or with pJC1-empty vector as control. Cultures were inoculated in CGXII containing 154 mM acetate, 222 mM glucose and 25 µg/ml kanamycin.

In *C. glutamicum*, the pyruvate carboxylase (PCx) encoded by *pyc* is the dominating HCO_3_^-^-dependent anaplerotic enzyme required for refueling the TCA cycle via oxaloacetate during growth on glycolytic carbon sources (24). In the following, we deleted *pyc* in the Δ*aceE* background to examine the influence of different CO_2_/HCO_3_^-^ levels on bicarbonate-dependent anaplerosis. Remarkably, the additional deletion of the *pyc* gene resulted in an elongated lag phase of about 20 hours and a reduced growth rate of strain Δ*aceE* Δ*pyc* (µ_max=_ 0.27 ± 0.02 h^-1^ compared to 0.40 ± h^-1^ of strain Δ*aceE*) during cultivation in CGXII minimal medium containing glucose and acetate (Fig. 2A). An extended lag phase was also observed for the Δ*pyc* single mutant, but the lag phase was significantly less prominent compared to the PDHC-deficient variant Δ*aceE* Δ*pyc* (Fig. 2C). The growth defect of both strains was successfully complemented by re-introducing the *pyc* gene on the plasmid pJC1 under control of its own promoters (Fig. 2 C). Remarkably, this extended lag phase was only observed in the presence of glucose, but not during cultivation on minimal medium containing acetate as sole carbon source (Fig. 2A).

### Glucose uptake results in strongly retarded growth in strains lacking pyruvate carboxylase

In order to verify whether glucose uptake caused the retarded growth, the Δ*aceE* Δ*pyc* strain was cultivated in the presence of 154 mM acetate and increasing amounts of glucose (100-250 mM). In line with our hypothesis, the lag phase showed a stepwise increase when higher amounts of glucose were added to the medium (Fig. 3A). Consistently, Δ*aceE* Δ*pyc* also showed significant elongated lag phases on the PTS sugars fructose and sucrose (Fig. S1). By contrast, the glucose concentration did not significantly affect the growth of the Δ*aceE* strain (Fig. 3A), in which the major anaplerotic enzyme PCx still replenishes the oxaloacetate pool.

**Figure 3:**
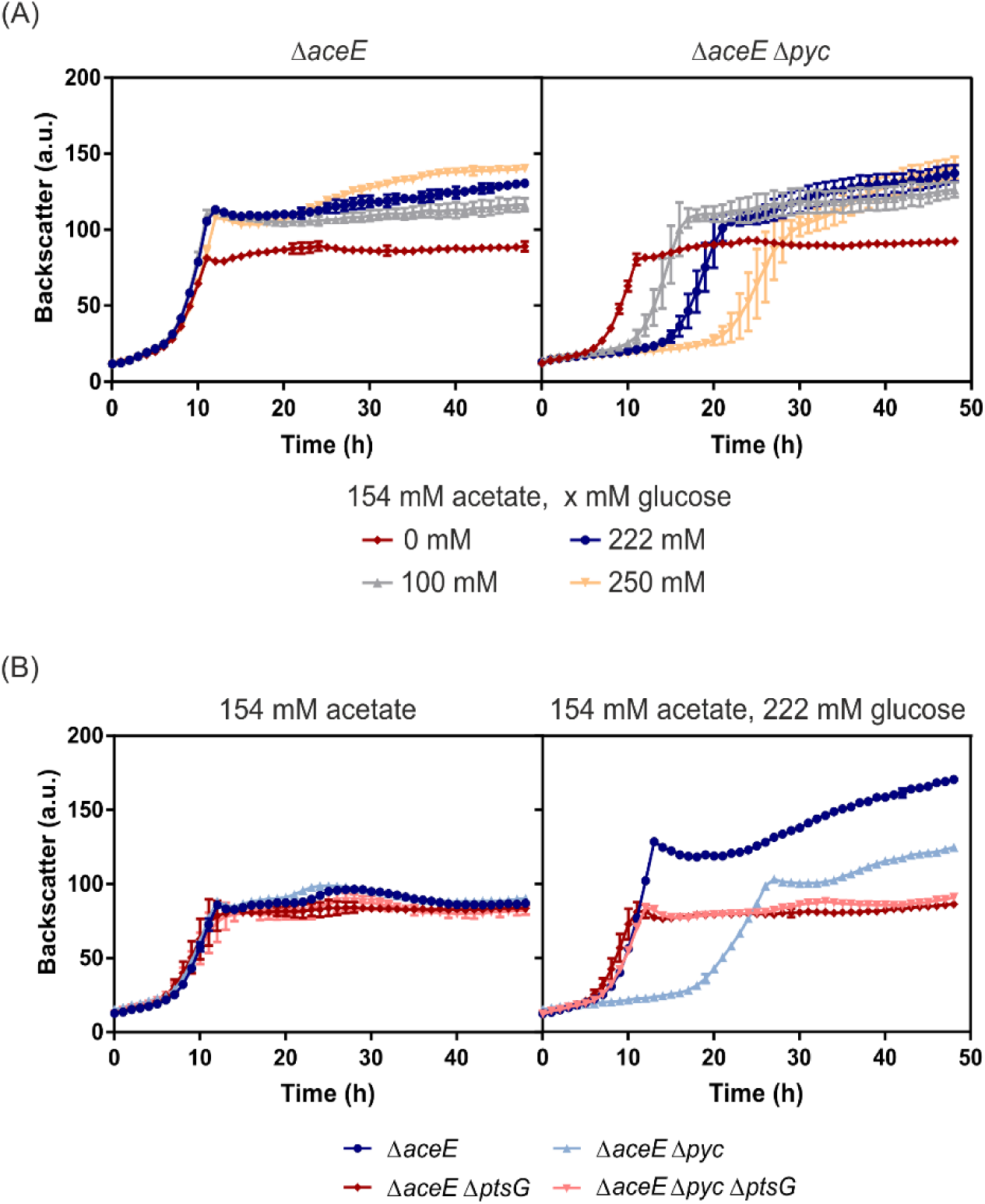
Influence of glucose consumption on the growth of *C. glutamicum* Δ*aceE* and Δ*aceE* Δ*pyc*. Growth curves shown are based on the backscatter measurements in a microtiter cultivation system, inoculated at an OD_600_ of 1. (A) The strains Δ*aceE* (left) and Δ*aceE* Δ*pyc* (right) were inoculated to an OD_600_ of 1 in CGXII medium containing 154 mM acetate and either 0 mM (red), 100 mM (grey), 222 mM (blue) or 250 mM glucose (orange). (B) Deletion of the *ptsG* gene in the Δ*aceE* and Δ*aceE* Δ*pyc* strains restored growth on glucose acetate mixtures (shades of red) when compared to the parental strains Δ*aceE* and Δ*aceE* Δ*pyc* (shades of blue). Data represent the average of three biological replicates, error bars the standard deviation.

To link the observed growth phenotype to the uptake of glucose, we deleted the *ptsG* gene, encoding the glucose specific permease of the PTS system, in the Δ*aceE* and Δ*aceE* Δ*pyc* strain background. Growth and glucose consumption rates of resulting strains were analyzed during growth on glucose and acetate. Remarkably, deletion of *ptsG* fully restored the growth of the parental strain Δ*aceE* Δ*pyc* (Δ*aceE* Δ*pyc*: 0.27 ± h^-1^; Δ*aceE* Δ*pyc* Δ*ptsG*: 0.39 ± 0.02 h^-1^), but resulted in reduced final backscatter values comparable to growth on acetate alone (Fig. 3B, Table 3). Strains lacking the *ptsG* gene consumed only minor amounts of glucose, while strains Δ*aceE* and Δ*aceE* Δ*pyc* consumed glucose after entering the stationary phase (Fig. S2). Glucose uptake rates of Δ*aceE* Δ*pyc* were significantly reduced to 10.87 ± 4.95 nmol min^-1^ g^-1^ compared to Δ*aceE* with 16.81 ± 3.01 nmol min^-1^ g^-1^. In contrast, deletion of *ptsG* did not restore growth on the PTS substrate fructose (Fig. S3), which is mainly imported by the permease encoded by *ptsF* (31, 32). Altogether, these findings clearly link the observed growth defect of the Δ*aceE* Δ*pyc* strain to the uptake of glucose or alternative PTS substrates.

**Table 1:**
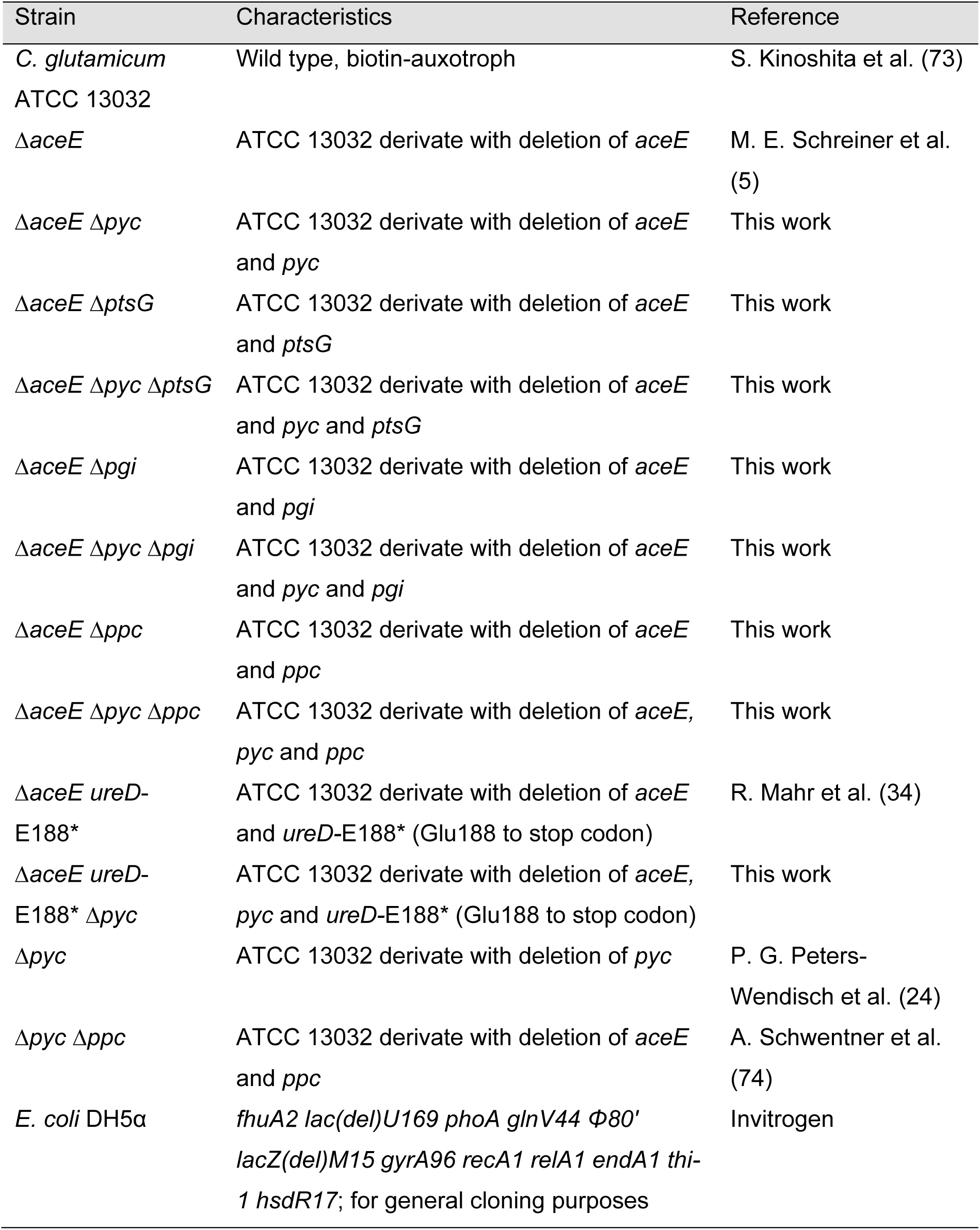
Bacterial strains used in this study.

**Table 2:**
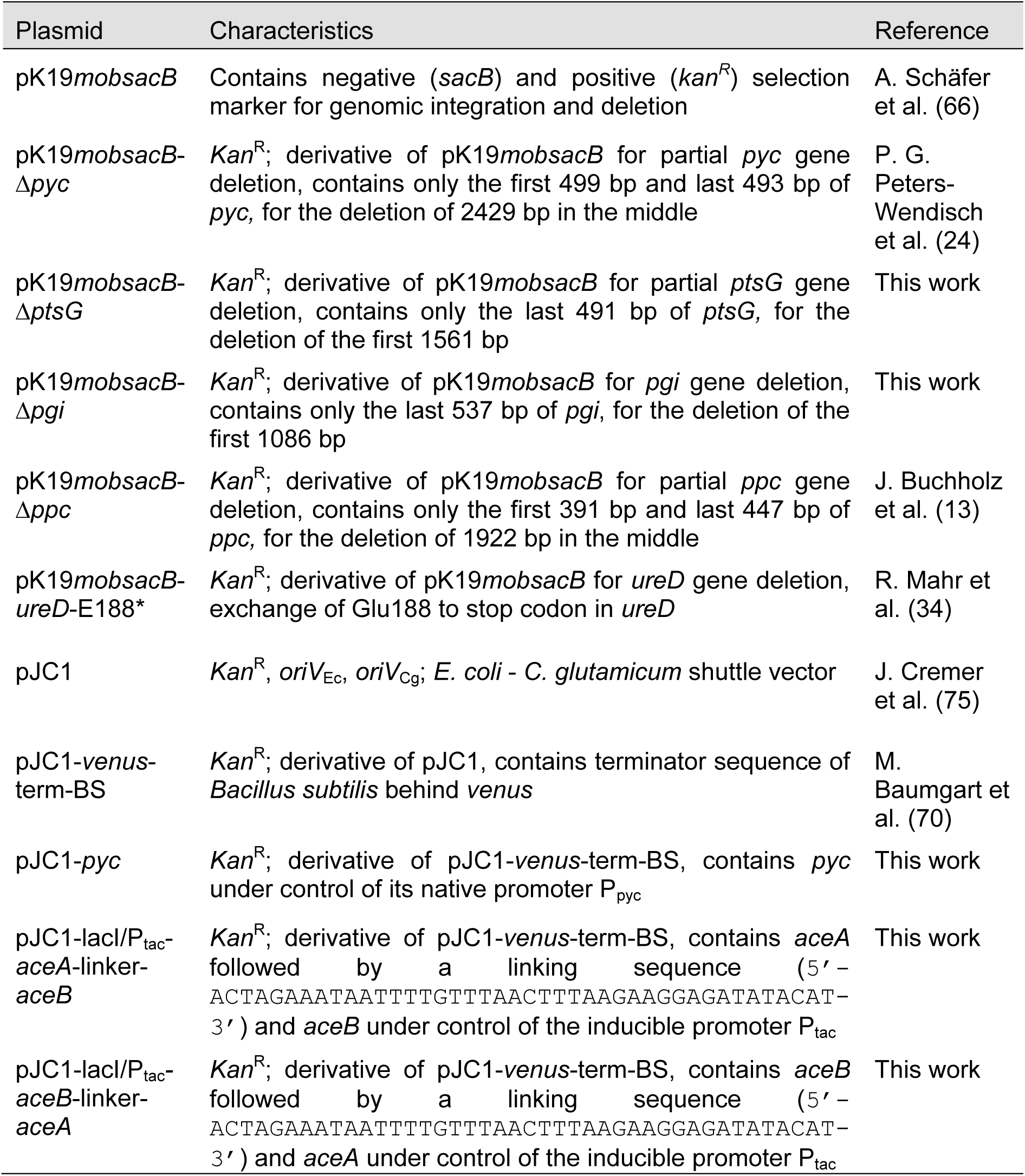
Plasmids used in this study.

**Table 3:** Glucose consumption rate of *C. glutamicum* PDHC-deficient strains in the exponential growth phase. For an overview of glucose consumption of the entire time course of the experiment including the stationary phase, the reader is referred to Fig. S2.

| Strain | Glucose consumption rate<br>(nmol min<sup>-1</sup> g<sup>-1</sup>)<sup>a</sup> |
| --- | --- |
| $\Delta aceE$ | 16.81 ± 3.01 |
| $\Delta aceE \Delta pyc$ | 10.87 ± 4.95 |
| $\Delta aceE \Delta ptsG$ | 7.24 ± 3.97 |
| $\Delta aceE \Delta pyc \Delta ptsG$ | 7.32 ± 1.88 |

<sup>a</sup> Means ± standard deviation of three independent biological replicates.

### Increased HCO_3_^-^ availability improved anaplerotic flux in *pyc* mutants

In the following, we used different strategies to alter the intracellular CO_2_/HCO_3_**^-^** pool. This was achieved by (1) supplementing the medium with HCO_3_**^-^**, (2) by mutation of the endogenous urease gene producing CO_2_ from urea in the early growth phase, (3) by increasing the pH of the growth medium, and (4) by re-routing metabolic flux via the pentose phosphate pathway (Fig. 3).

Similar as observed for the parental strain Δ*aceE*, addition of 100 mM HCO_3_^-^ eliminated the lag phase of PDHC-deficient strains lacking the *pyc* gene (Fig. 2B) during microtiter plate cultivation on glucose-acetate mixtures. Restoration of growth was not possible in a strain lacking both anaplerotic enzymes PCx and PEPCx, indicating that the positive effect of HCO_3_^-^ on the *ΔaceE Δpyc* strain depends on the increased flux over PEPCx (Fig. 4A, Fig. S4). In contrast to PCx-deficient strains, no negative impact of glucose on growth of the strain Δ*aceE* Δ*ppc* was observed (Fig. S5) confirming again the superior role of PCx in *C. glutamicum* (24, 33). Interestingly, we also observed a significant impact of the culture volume on growth of Δ*aceE* Δ*pyc.* Since efficient liquid-air mixing will continuously remove CO_2_ from the culture medium, the lag phase of Δ*aceE* Δ*pyc* was much more prominent during microtiter plate cultivation (∼17 h) compared to cultures in shake flasks (∼7 h lag phase in 100 ml cultures; ∼13 h lag phage in 50 ml) (Fig. S6).

**Figure 4:**
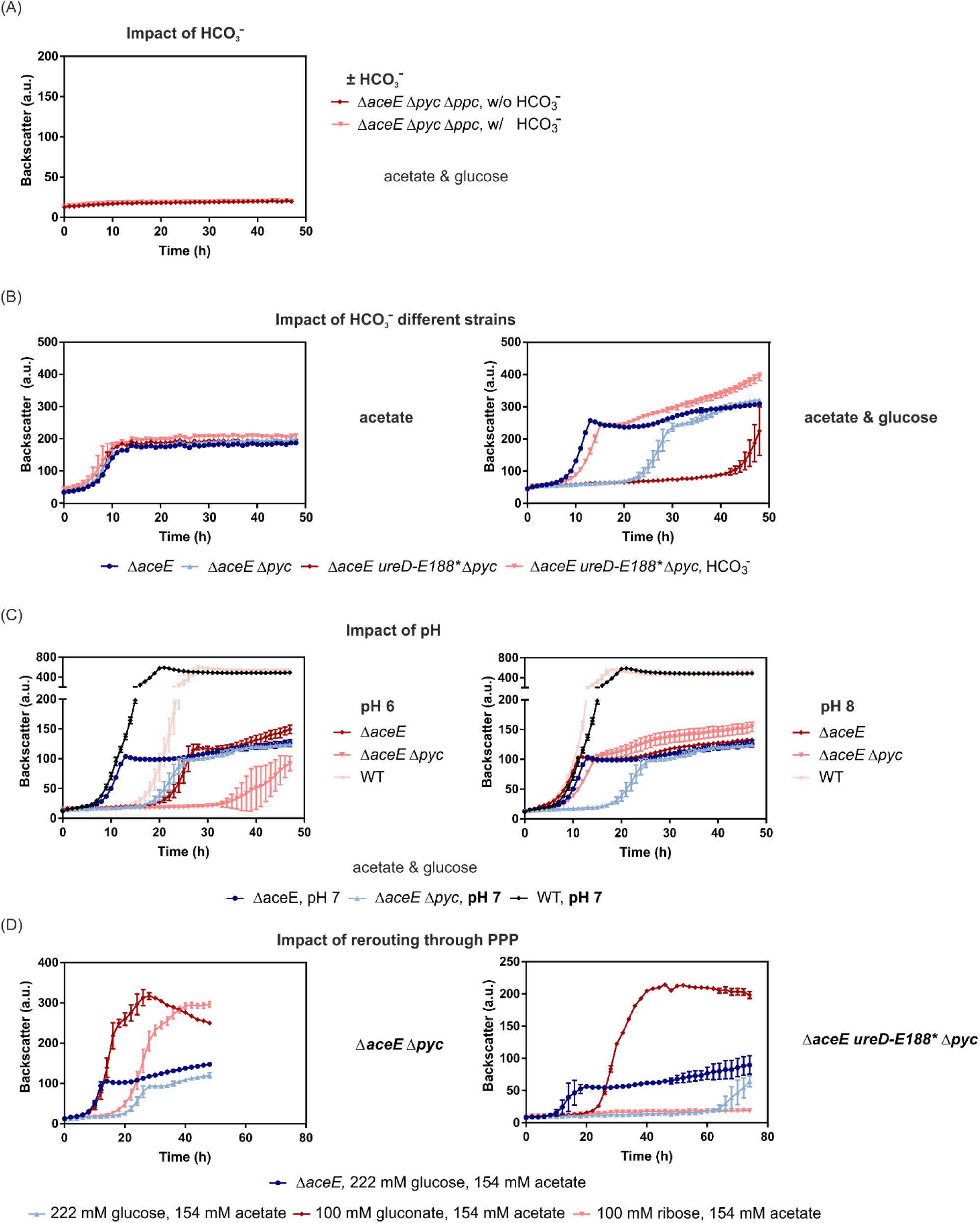
Increased availability of HCO_3_^-^ improved anaplerotic flux via PEPCx in *pyc*-deficient *C. glutamicum* strains. Growth curves shown are based on the backscatter measurements in a microtiter cultivation system, inoculated at an OD_600_ of 1. Symbols represent the backscatter means of biological triplicates (n=3). (A) Strain Δ*aceE* Δ*pyc* Δ*ppc* was inoculated in CGXII medium containing 154 mM acetate and 222 mM glucose (shown in dark red) and, if indicated, 100 mM KHCO_3_^-^ was added (shown in light red). (B) The strains Δ*aceE* and Δ*aceE* Δ*pyc* as well as the WT were cultivated in CGXII medium containing 154 mM acetate and 222 mM glucose adjusted to a pH of 6 (left) or pH of 8 (right) in comparison to cultivation at pH 7. (C) The strains Δ*aceE* and Δ*aceE ureD*-E188* Δ*pyc* were cultivated in CGXII media containing 154 mM acetate (left) or 154 mM acetate and 222 mM glucose (right) with and without the addition of KHCO_3_^-^. (D) The strains Δ*aceE,* Δ*aceE* Δ*pyc* (left), Δ*aceE ureD*-E188* Δ*pyc* (right) were inoculated in different growth media containing 154 mM acetate and 222 mM glucose (shades of blue), 100 mM gluconate (red) or 100 mM ribose (light red).

The urease enzyme, catalyzing the degradation of urea to ammonium and carbon dioxide, represents a further important contributor to the intracellular CO_2_/HCO_3_**^-^** pool, especially in the early exponential phase when cell densities and decarboxylation reaction rates are low. In a previous study, the *ureD*-E188* mutation was found to abolish urease activity in *C. glutamicum* and to enhance L-valine production of PDHC-deficient strains (34). In this study, we introduced the *ureD*-E188* mutation into the Δ*aceE* Δ*pyc* background to examine the influence of an even lower CO_2_/HCO_3_^-^ level on anaplerosis in this particular strain background. The additional inactivation of the urease enzyme resulted in a significantly elongated lag phase of about 46 hours, which could also be complemented by addition of HCO_3_^-^ (Fig. 4B) demonstrating the importance of the CO_2_/HCO_3_^-^ pool during the initial growth phase. Furthermore, we assessed the impact of urease mutation on L-valine and L-alanine production of strain Δ*aceE* and strain Δ*aceE ureD*-E188*and the complementation via addition of HCO_3_^-^. Within 28 hours of cultivation, the Δ*aceE* strain produced 15 mM L-valine and 10 mM L-alanine as by-product, while the Δ*aceE ureD*-E188* accumulated 54 mM L-valine and 38 mM L-alanine (Fig. 5). Addition of HCO_3_^-^, however, significantly reduced L-valine production by 54% (Δ*aceE*) and 59% (Δ*aceE ureD*-E188*), respectively. The by-product L-alanine increased by 22% for the Δ*aceE* strain and decreased by 51% for the Δ*aceE ureD*-E188* strain (Fig. 5). The example of L-valine/L-alanine production illustrates the important role of the intracellular CO_2_/HCO_3_^-^ levels on metabolic flux in PDHC-deficient strains.

**Figure 5:**
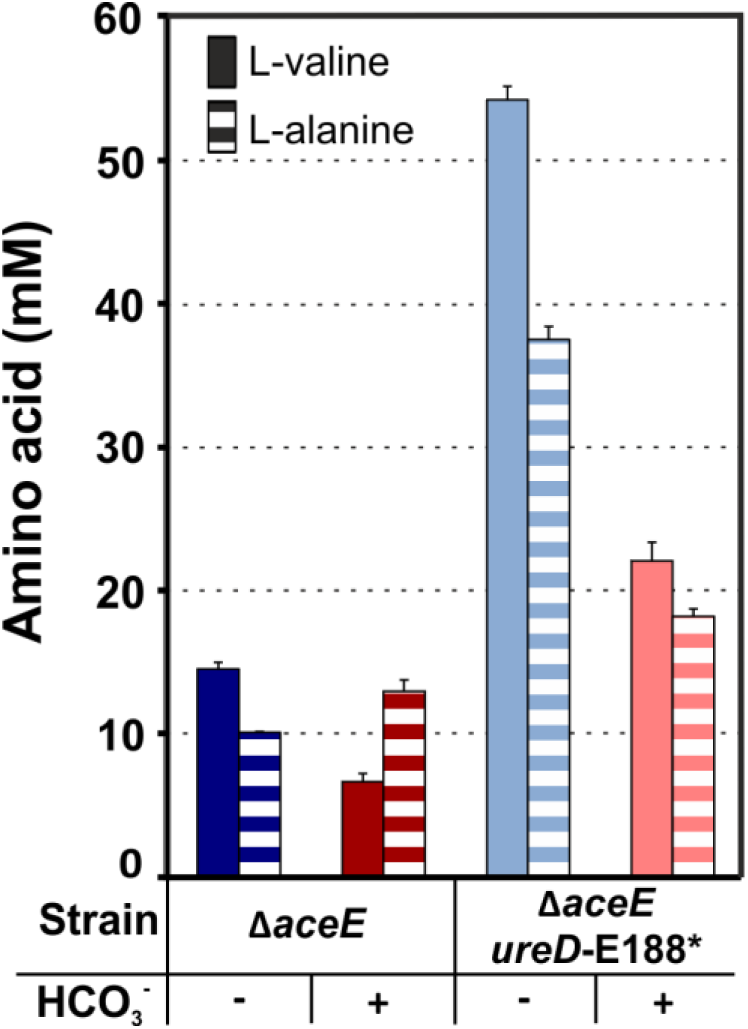
The intracellular CO_2_/HCO_3_^-^ pool significantly affects L-valine production. Investigations on the influence of carbonate on growth and L-valine production of *C. glutamicum* Δ*aceE* and Δ*aceE ureD*-E188* are shown. L-valine (filled bars) and L-alanine (dashed bars) titers in the supernatant after 28 h of cultivation in the microtiter cultivation system. Data represent average values from three independent biological replicates using ultra-high performance liquid chromatography (uHPLC).

Another approach to influence the extracellular HCO_3_^-^ availability was to change the pH conditions (2). According to the Bjerrum plot, CO_2_ is predominant at acidic conditions, while at alkaline conditions CO_3_^2-^ is the dominant form. At a pH of about 8, HCO_3_^-^ is the prevalent form. As HCO_3_^-^ is the substrate of the anaplerotic enzymes PCx and PEPCx, the pH in the cultivation medium was adjusted to pH 8 using KOH leading to an equilibrium shift towards higher HCO_3_^-^ availability. As negative control, the pH was adjusted to pH 6 using HCl. While growth of both strains was retarded at pH 6, it was significantly enhanced at pH 8 (Fig. 4C). Thus, elevation of the culture pH improved growth of PDHC-deficient strains likely by increasing anaplerotic flux. With the reaction catalyzed by the 6-phosphogluconate dehydrogenase (6PGDH), the PPP comprises another decarboxylation reaction in central carbon metabolism, thereby contributing to the intracellular CO_2_/HCO_3_**^-^**pool. Remarkably, the lag phases of Δ*aceE* Δ*pyc* and Δ*aceE ureD-*E188* Δ*pyc* were mostly complemented during growth on acetate and gluconate – the latter entering the PPP *via* gluconate-6-phosphate (Fig. 4D). Growth on ribose, which enters the PPP via ribulose-5-phosphate and thereby bypassing the decarboxylation catalyzed by 6PGDH, showed a significantly elongated lag phase in *pyc* deficient strains (Fig. 4D). In a further attempt, metabolic/glycolytic flux was re-routed through the PPP by deleting *pgi*, encoding the glucose-6-phosphate isomerase. However, deletion of *pgi* resulted in a severe growth defect in the Δ*aceE* background. In this context the growth of Δ*aceE* Δ*pyc* featured only a minor, non-significant improvement (Fig. S7).

Taken together, our findings highlight the important impact of the intracellular CO_2_/HCO_3_^-^ pool on metabolic flux in central carbon metabolism, which is especially evident in strain Δ*aceE* Δ*pyc* lacking a central decarboxylation reaction and the key carboxylase PCx in *C. glutamicum*.

### Refueling the TCA cycle improves growth of *pyc* mutants

Glucose catabolism requires sufficient anaplerotic flux to replenish TCA cycle intermediates providing precursors for various anabolic pathways. Therefore, we tested whether the addition of TCA intermediates would complement the negative impact of glucose on the growth of Δ*aceE* Δ*pyc*. Remarkably, all tested TCA intermediates (succinate, malate, citrate and the glutamate containing dipeptide Glu-Ala) reduced the extended lag phase of Δ*aceE* Δ*pyc* during growth on glucose-acetate mixtures (Fig. 6A). The dipeptide Glu-Ala and succinate reduced the elongated lag phase to a high extent by 90 % (Δt = 10 %) and 85 % (Δt = 15 %), respectively, while the effects of citrate and malate were weaker (Δt = 40 % and 38 %, respectively) (Table 4).

**Figure 6:**
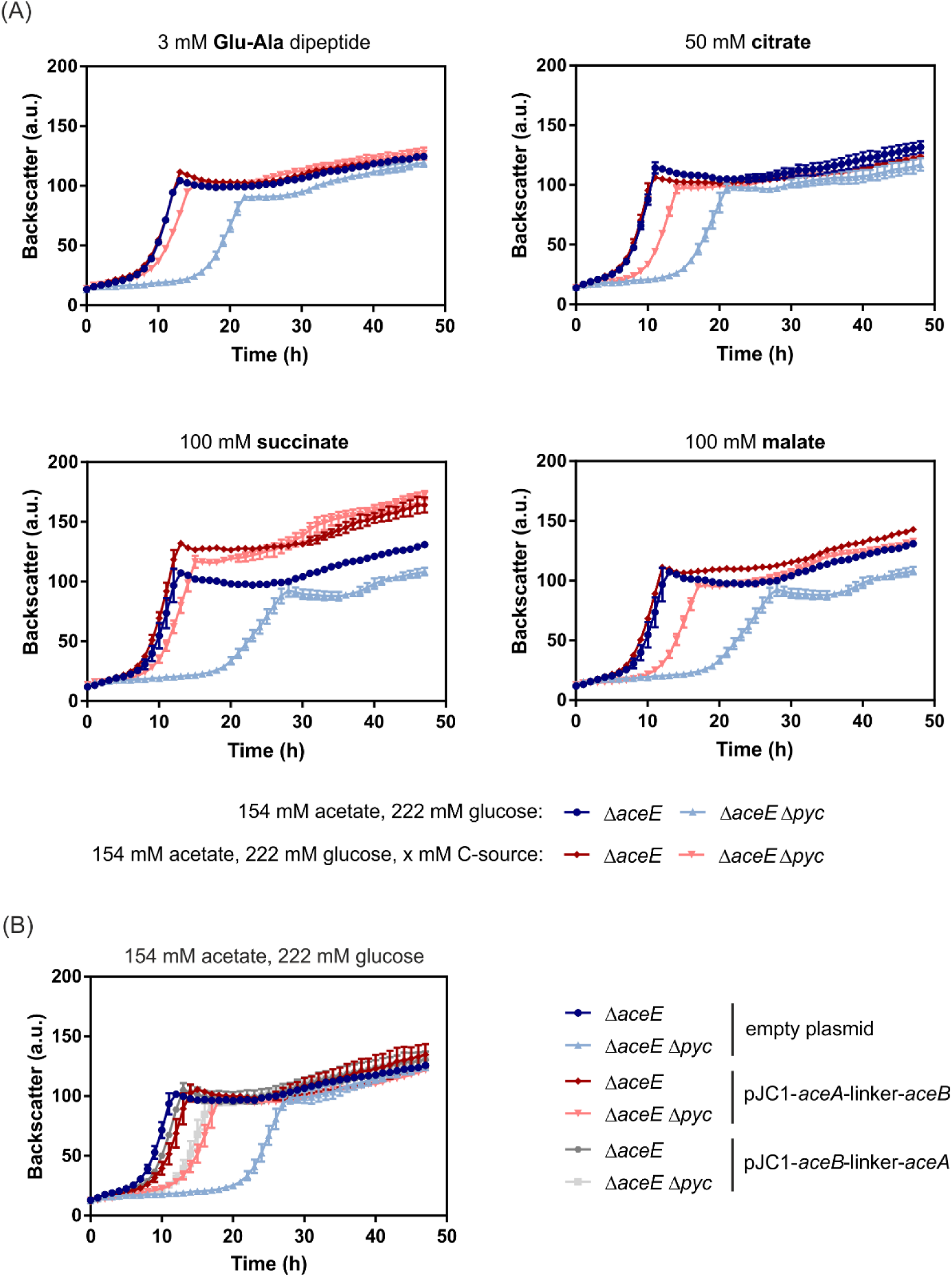
Refueling the TCA cycle to complement glucose intolerance of Δ*aceE* Δ*pyc*. Growth curves shown are based on the backscatter measurements in a microtiter cultivation system, while symbols represent the backscatter means of biological triplicates (n=3). Strains Δ*aceE* and Δ*aceE* Δ*pyc* were inoculated to an OD_600_ of 1 in CGXII medium with 154 mM acetate and 222 mM glucose. (A) The TCA fueling carbon sources glutamate (3 mM Glu-Ala dipeptide), citrate (50 mM), succinate (100 mM) or malate (100 mM) were added to the medium to analyze their effect on lag phase complementation (shades of red). As a control, growth on 154 mM acetate and 222 mM glucose is shown for Δ*aceE* (dark blue) and Δ*aceE* Δ*pyc* (light blue) in each experiment. (B) Strains Δ*aceE* and Δ*aceE* Δ*pyc* were transformed with pJC1-*aceA*-linker-*aceB* or pJC1-*aceB*-linker-*aceA* plasmid for simultaneous overexpression of the glyoxylate shunt enzymes isocitrate lyase (*aceA*) and malate synthase (*aceB*), while transformation of Δ*aceE* Δ*pyc* and Δ*pyc* with pJC1-empty vector served as control. Cultures were inoculated to an OD_600_ of 1 in CGXII containing 154 mM acetate, 222 mM glucose and 25 μg/ml kanamycin.

**Table 4:** Overview of lag phases and growth rates of *C. glutamicum* Δ*aceE* and Δ*aceE* Δ*pyc* cultivated on different TCA cycle carbon sources in CGXII medium containing 154 mM acetate and 222 mM glucose.

| Added carbon source* | $\Delta aceE$ | | | $\Delta aceE \Delta pyc$ | | |
| --- | --- | --- | --- | --- | --- | --- |
| | Lag phase | $\Delta t$ (%)** | Growth rate ($h^{-1}$) | Lag phase | $\Delta t$ (%)** | Growth rate ($h^{-1}$) |
| / | - | 0 | $0.41 \pm 0.01$ | + | 100 | $0.27 \pm 0.02$ |
| Glu-Ala (3 mM) | - | 0 | $0.35 \pm 0.01$ | ÷ | 10 | $0.33 \pm 0.01$ |
| Citrate (50 mM) | - | 0 | $0.39 \pm 0.01$ | ÷ | 40 | $0.35 \pm 0.01$ |
| Succinate (100 mM) | - | -8 | $0.36 \pm 0.01$ | ÷ | 15 | $0.34 \pm 0.02$ |
| Malate (100 mM) | - | -8 | $0.35 \pm 0.01$ | ÷ | 38 | $0.35 \pm 0.02$ |

\*Additionally the medium contained 154 mM acetate and 222 mM glucose,
\*\* The difference between the first doubling time of strain $\Delta aceE$ and $\Delta aceE \Delta pyc$ was defined as 100%. + = elongated lag phase referring to growth on 154 mM acetate with 222 mM glucose, ÷ = shorter lag phase, - = no lag phase.

Based on previous studies, it was known that the glyoxylate shunt might be switched off during growth on glucose-acetate mixtures, due to the inhibitory effect of glucose (35). Here, overexpression of the genes *aceA* and *aceB,* encoding the glyoxylate shunt enzymes isocitrate lyase and malate synthase, resulted in a significantly shorter lag phase of strain Δ*aceE* Δ*pyc* (Fig. 6B). Overexpression was accomplished using the vector pJC1 harboring an inducible P*_tac_* promoter in front of two synthetic operon variants: pJC1-*aceA*-linker-*aceB* and pJC1-*aceB*-linker-*aceA*. However, it has to be noted that the overexpression of *aceA* and *aceB* led to slightly longer lag phases of strain Δ*aceE* than compared to the empty vector control (Fig. 6B). Expression from the leaky P*_tac_* promoter yielded the best result, while induction with IPTG resulted in severe growth defects (data not shown).

Unusual intermediate accumulation or depletion can provide valuable information regarding intracellular flux imbalances causing the observed growth defects. Gas chromatography-time of flight (GC-ToF) analysis was performed for analysis of the metabolic states by comparing samples of Δ*aceE* during early exponential phase, Δ*aceE* Δ*pyc* at the same time point during the lag phase, as well as Δ*aceE* Δ*pyc* during early exponential phase (see Fig. S8 showing the sampling scheme). Additionally, samples of both PDHC-deficient strains cultivated with 100 mM HCO_3_^-^ were measured and WT cells during exponential phase were used as reference (Table S3). No significant intermediate accumulation or occurrence of unusual compounds was observed in the lag phase sample of Δ*aceE* Δ*pyc* compared to all other samples. Although oxaloacetate cannot be measured by this method, a complete depletion of oxaloacetate-derived aspartate was detected in Δ*aceE* Δ*pyc* cells during the lag phase but not during exponential phase or in HCO_3_^-^ complemented samples. This supports the hypothesis that oxaloacetate depletion is the key reason for the elongated lag phase.

In order to identify oxaloacetate depletion as key growth limiting factor and to decouple this effect from anaplerosis-dependent replenishment, strains Δ*pyc*, Δ*pyc* Δ*ppc* and Δ*aceE* Δ*pyc* Δ*ppc* were tested in the presence of different TCA intermediates. Here, mutants lacking both anaplerotic reactions (PCx and PEPCx, strains Δ*pyc* Δ*ppc* and Δ*aceE* Δ*pyc* Δ*ppc*) were not able to grow on glucose acetate mixtures at all, but grew on gluconeogenetic substrates like acetate (Fig. 4A, Fig. S4). Again, all TCA intermediates were able to complement the glucose sensitivity of these *pyc*-deficient strains and the growth of Δ*pyc* Δ*ppc* and Δ*aceE* Δ*pyc* Δ*ppc* was effectively restored (Table 5). Nevertheless, slight differences were observed, which might also be caused by differences in the uptake of these carbon sources.

**Table 5:** Overview of lag phases and growth rates of *C. glutamicum* Δ*pyc*, Δ*pyc* Δ*ppc* and Δ*aceE* Δ*pyc* Δ*ppc* cultivated on different TCA cycle carbon sources in CGXII medium containing 154 mM acetate and 222 mM glucose.

| Added carbon source* | $\Delta pyc$ | | $\Delta pyc \Delta ppc$ | | $\Delta aceE \Delta pyc \Delta ppc$ | |
| --- | --- | --- | --- | --- | --- | --- |
| | $\Delta t_{lag}^{**}$ | Growth rate ($h^{-1}$)<sup>***</sup> | $\Delta t_{lag}^{**}$ | Growth rate ($h^{-1}$)<sup>***</sup> | $\Delta t_{lag}^{**}$ | Growth rate ($h^{-1}$)<sup>***</sup> |
| / | 6 h | $0.17 \pm 0.02$ | No growth | | No growth | |
| Glu-Ala (3 mM) | n.d. | n.d. | 8 h | $0.21 \pm 0.01$ | 30 h | $0.14 \pm 0.01$ |
| Citrate (50 mM) | 3 h | $0.19 \pm 0.01$ | 5 h | $0.26 \pm 0.01$ | 8 h | $0.20 \pm 0.03$ |
| Succinate (100 mM) | 2 h | $0.2 \pm 0.02$ | 3 h | $0.16 \pm 0.03$ | 5 h | $0.12 \pm 0.03$ |
| Malate (100 mM) | 3 h | $0.21 \pm 0.01$ | 17 h | $0.18 \pm 0.01$ | 18 h | $0.24 \pm 0.01$ |

\*Additionally the medium contained 154 mM acetate and 222 mM glucose,
\*\* = $\Delta t_{lag}$ represents the difference in lag phase duration (h) between $\Delta aceE$ growing on glucose-acetate mixture and the strain of interest growing on the respective carbon source(s).
\*\*\* = for comparison, growth rate in glucose-acetate mixture was $0.41 h^{-1}$ for $\Delta aceE$ and $0.28 h^{-1}$ for $\Delta aceE \Delta pyc$ in this experiment. n.d. = not determined

### Adaptation of *C. glutamicum* Δ*aceE* Δ*pyc* to growth on glucose and acetate

In a previous study, Kotte et. al reported on an elongated lag phase of *E. coli* as a result of glucose-gluconeogenetic substrate shifts (36). This phenomenon was ascribed to the formation of a small subpopulation, which is able to start growing after carbon source switches (36). Usually, the growth of a small subpopulation is obscured by typical bulk measurements, but can be visualized by single-cell approaches, such as flow cytometry. To this end, the membranes of Δ*aceE* and Δ*aceE* Δ*pyc* cells were stained using the non-toxic, green fluorescent dye PKH67 prior to inoculation. By cellular growth, the amount of fluorescent dye is diluted by membrane synthesis leading to a decrease of single cell PKH67 fluorescence (36). Stained Δ*aceE* and Δ*aceE* Δ*pyc* cells were cultivated in minimal medium containing glucose and acetate (Fig. 7A). As non-proliferating control, Δ*aceE* Δ*pyc* cells were incubated in glucose as sole carbon source. Using flow cytometry, membrane staining was analyzed during the cultivation at the single-cell level (Fig. 7B). While the mean of fluorescence of the whole Δ*aceE* population decreased from 2.7 × 10^4^ a.u. to 7.5 × 10^2^ a.u., the mean of the Δ*aceE* Δ*pyc* population incubated in glucose medium only shifted from 3.1 × 10^4^ a.u. to 1.3 × 10^3^ a.u. resulting from a minor dilution/bleaching of fluorescence intensity during incubation. Remarkably, both strains, Δ*aceE* and Δ*aceE* Δ*pyc,* featured a rather heterogeneous adaptation behavior on glucose and acetate mixtures which was apparently time delayed in strain Δ*aceE* Δ*pyc* reflecting the elongated lag phase. Already in the early phase of cultivation, the populations split up in two subpopulations of low and high PKH67 fluorescence indicating that only a few cells start to proliferate, while a fraction of the population was not able to grow under same conditions (37–39).

**Figure 7:**
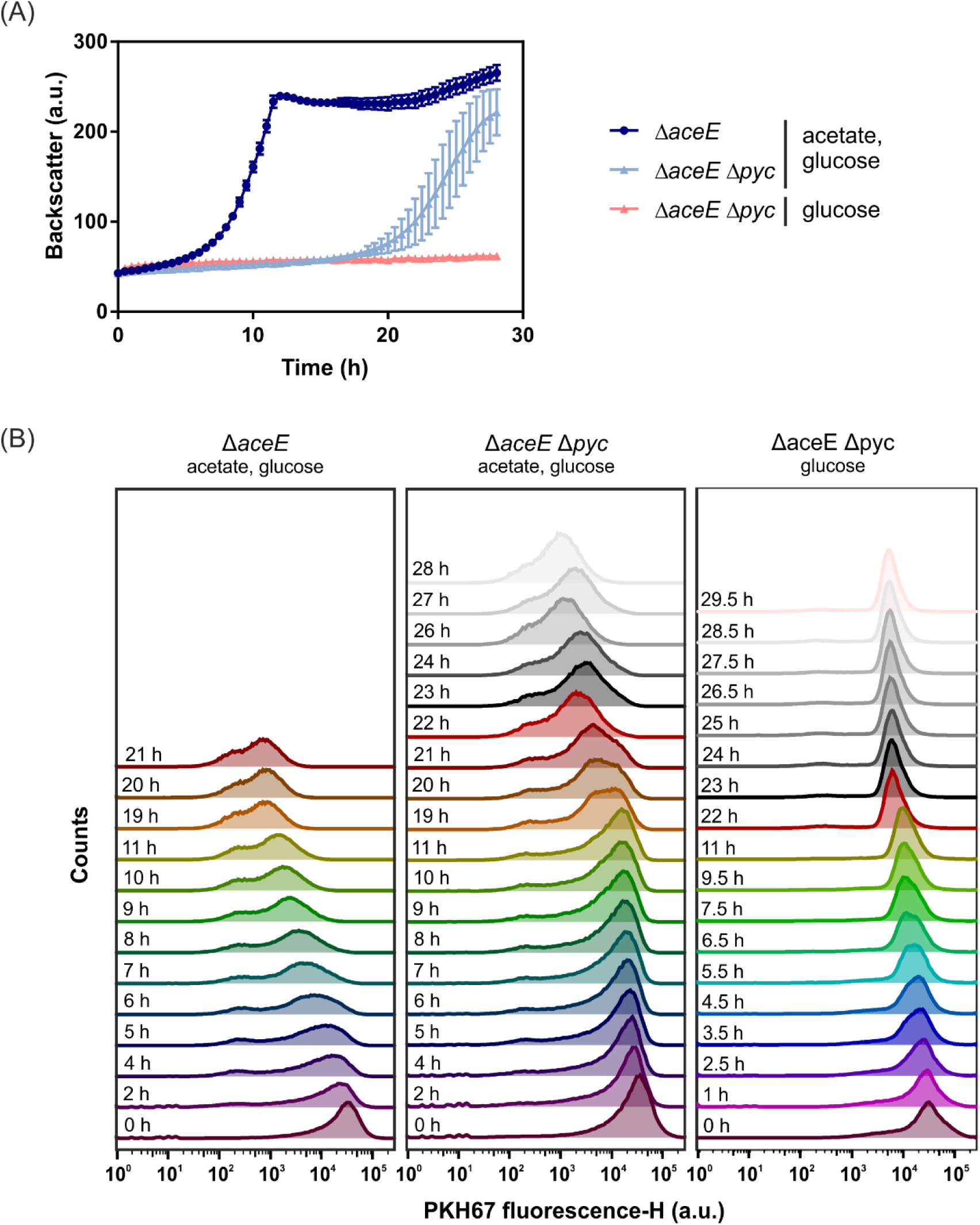
Heterogeneous adaptation of *C. glutamicum* Δ*aceE* and Δ*aceE* Δ*pyc* during growth on glucose and acetate. To identify a potential growing subpopulation the fluorescent dye PKH67 was used for membrane staining of Δ*aceE* and Δ*aceE* Δ*pyc* prior to cultivation. Cells were cultivated in a microtiter cultivation system in the presence of 154 mM acetate and 222 mM glucose. As non-growing control, Δ*aceE* Δ*pyc* was additionally incubated in CGXII containing solely 222 mM glucose. Cell growth (A) was online monitored using a microtiter cultivation system and staining intensities were measured on single cell level during cultivation using flow cytometry (n=3 biological replicates). (B) Shown is the frequency of PKH67 staining intensities distributions of one representative culture of each sample over the time course of cultivation (A).

To study the adaptation of Δ*aceE* Δ*pyc*, comparative transcriptome analysis were performed using DNA microarrays. To this end, Δ*aceE* and Δ*aceE* Δ*pyc* cells were harvested at a comparable optical density during early exponential phase (Fig. S8). Both strains were cultivated in CGXII media with 154 mM acetate and 222 mM glucose. Under the chosen conditions, a total of 354 genes showed a significantly altered mRNA level. While 121 genes were at least 1.7-fold upregulated (p-value < 0.05), 233 genes were 0.7-fold downregulated (p-value < 0.05) in strain Δ*aceE* Δ*pyc* (Table S4). An overview of expression changes of selected genes is shown in Table 6. Among the various changes on the level of gene expression, we observed a 0.59-fold and 0.67-fold downregulation of the respective PTS-system components *ptsI* and *ptsG* in strain Δ*aceE* Δ*pyc*, which is in line with the decreased glucose uptake rates of this strain (Table 3). In contrast, *iolT1* encoding the *myo*-inositol transporter, which is inter alia responsible for PTS-independent glucose uptake, was upregulated (40, 41). This is in line with the fact that the *ptsG* deficient strains still consumed minor amounts of glucose. Interestingly, genes involved in acetate metabolism, including *aceA* and *aceB* constituting the glyoxylate shunt, displayed an increased mRNA level in strain Δ*aceE* Δ*pyc*. This may also represent an important adaptive mechanism, as the glyoxylate shunt needs to be active in Δ*aceE* Δ*pyc* to compensate for the loss of TCA intermediates during co-consumption of acetate and glucose as demonstrated by the effect of *aceA*-*aceB* overexpression in this study (Fig. 6B). While the mRNA level of genes encoding anaplerotic enzymes did not show a significant change, the glycolytic genes *pgi* and *pyk* featured an about two-fold reduced level in the Δ*aceE* Δ*pyc* mutant. This might as well reflect an adaptive mechanism, as a downregulation of *pyk* would reduce flux from PEP to pyruvate and reduced expression of *pgi* would probably lead to an increased flux through the PPP contributing an additional decarboxylation step.

**Table 6:**
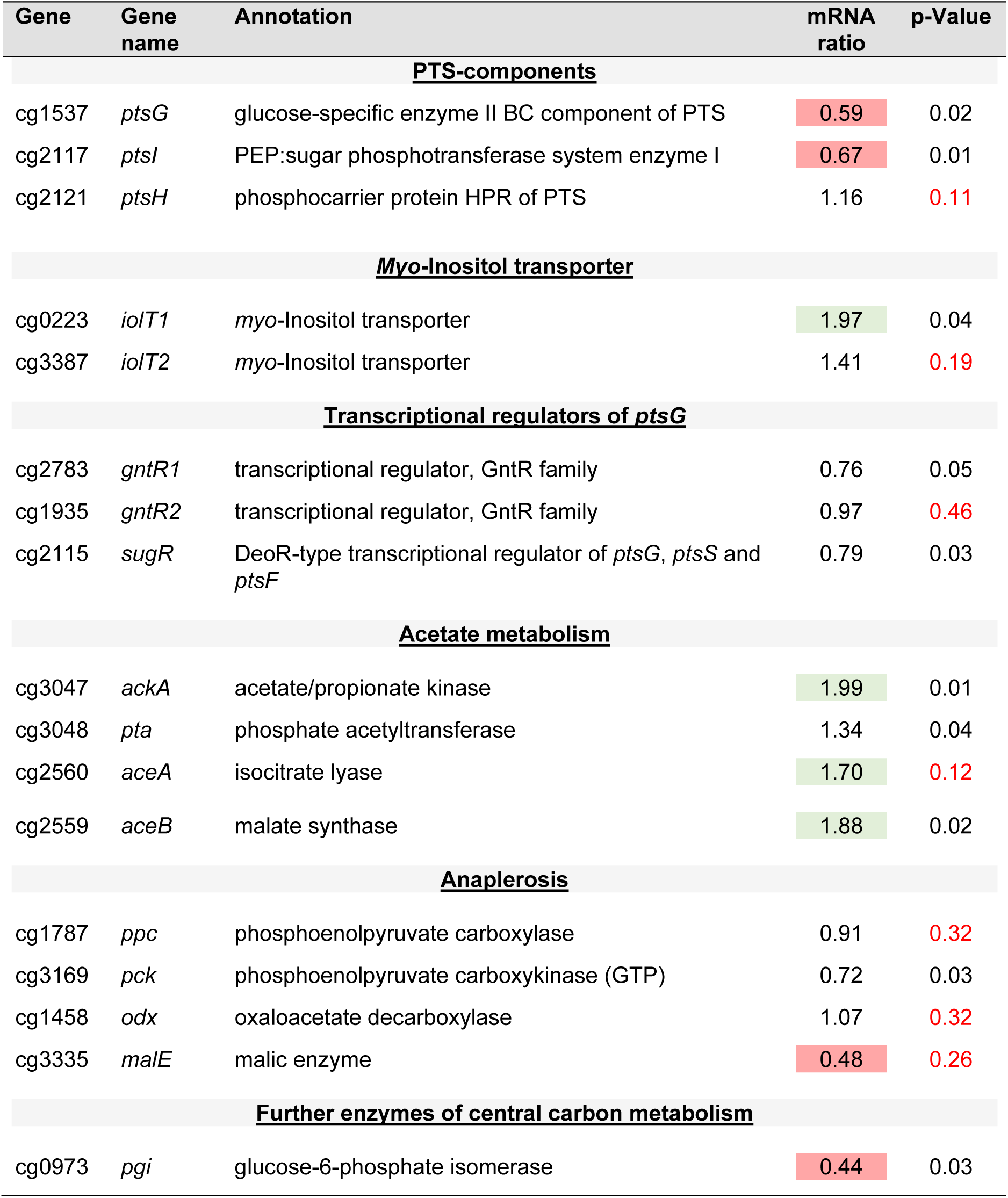
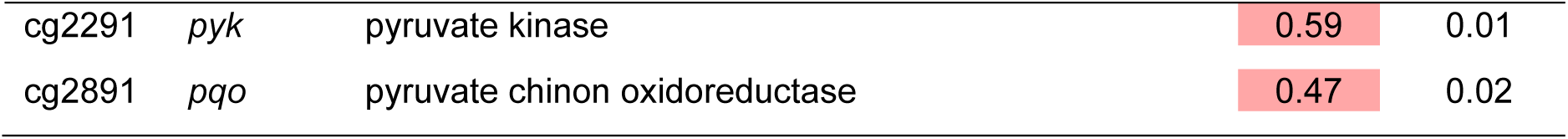
Comparative transcriptome analysis of *C. glutamicum* Δ*aceE* Δ*pyc* and strain Δ*aceE* and during growth on glucose and acetate. Shown are selected genes encoding for central metabolic enzymes or regulators (for mRNA ratios of Δ*aceE* Δ*pyc* versus Δ*aceE* ≤ 0.70 values are shaded in red; mRNA ratio ≥ 1.70 are shaded in green). p-values >0.05 are highlighted red (n=3 biological replicates). For the complete dataset, the reader is referred to Table S3.

| Gene | Gene name | Annotation | mRNA ratio | p-Value |
| --- | --- | --- | --- | --- |
| **<u>PTS-components</u>** |  |  |  |  |
| cg1537 | *ptsG* | glucose-specific enzyme II BC component of PTS | 0.59 | 0.02 |
| cg2117 | *ptsI* | PEP:sugar phosphotransferase system enzyme I | 0.67 | 0.01 |
| cg2121 | *ptsH* | phosphocarrier protein HPR of PTS | 1.16 | 0.11 |
| **<u>Myo-Inositol transporter</u>** |  |  |  |  |
| cg0223 | *iolT1* | myo-Inositol transporter | 1.97 | 0.04 |
| cg3387 | *iolT2* | myo-Inositol transporter | 1.41 | 0.19 |
| **<u>Transcriptional regulators of *ptsG*</u>** |  |  |  |  |
| cg2783 | *gntR1* | transcriptional regulator, GntR family | 0.76 | 0.05 |
| cg1935 | *gntR2* | transcriptional regulator, GntR family | 0.97 | 0.46 |
| cg2115 | *sugR* | DeoR-type transcriptional regulator of *ptsG*, *ptsS* and *ptsF* | 0.79 | 0.03 |
| **<u>Acetate metabolism</u>** |  |  |  |  |
| cg3047 | *ackA* | acetate/propionate kinase | 1.99 | 0.01 |
| cg3048 | *pta* | phosphate acetyltransferase | 1.34 | 0.04 |
| cg2560 | *aceA* | isocitrate lyase | 1.70 | 0.12 |
| cg2559 | *aceB* | malate synthase | 1.88 | 0.02 |
| **<u>Anaplerosis</u>** |  |  |  |  |
| cg1787 | *ppc* | phosphoenolpyruvate carboxylase | 0.91 | 0.32 |
| cg3169 | *pck* | phosphoenolpyruvate carboxykinase (GTP) | 0.72 | 0.03 |
| cg1458 | *odx* | oxaloacetate decarboxylase | 1.07 | 0.32 |
| cg3335 | *malE* | malic enzyme | 0.48 | 0.26 |
| **<u>Further enzymes of central carbon metabolism</u>** |  |  |  |  |
| cg0973 | *pgi* | glucose-6-phosphate isomerase | 0.44 | 0.03 |
| cg2291 | *pyk* | pyruvate kinase | 0.59 | 0.01 |
| cg2891 | *pqo* | pyruvate chinon oxidoreductase | 0.47 | 0.02 |

### Less is more - Adaptive laboratory evolution of Δ*aceE* Δ*pyc* revealed rapid inactivation of glucose uptake

In order to identify key mutations abolishing the lag phase of Δ*aceE* Δ*pyc*, an adaptive laboratory evolution (ALE) experiment was performed. In this ALE, *C. glutamicum* Δ*aceE* Δ*pyc* was grown in CGXII minimal medium containing 154 mM acetate and 222 mM glucose in overall 16 repetitive-batch cultures. Remarkably, already after eight inoculations a significantly shortened lag phase was observed; after 10 inoculations strain Δ*aceE* Δ*pyc* featured a similar lag phase as the strain Δ*aceE* (Fig. 8A). In contrast, serial cultivation transfers in CGXII medium containing solely acetate did not lead to improved growth in glucose-acetate medium indicating that glucose acted as evolutionary selection pressure (Fig. S9). Genome re-sequencing of three selected glucose-acetate clones presented in Fig. 8B revealed mutations in the *ptsI* gene, encoding the EI enzyme of the PTS system, in two independently evolved clones (Table 7). This is in agreement with the reduced biomass formation of both clones, which apparently abolished glucose uptake to optimize growth on acetate. The third clone showed only a slightly accelerated growth, but was not impaired in biomass formation. Here, sequencing revealed a mutation in the *rpsC* gene, encoding the ribosomal S3 protein.

**Figure 8:**
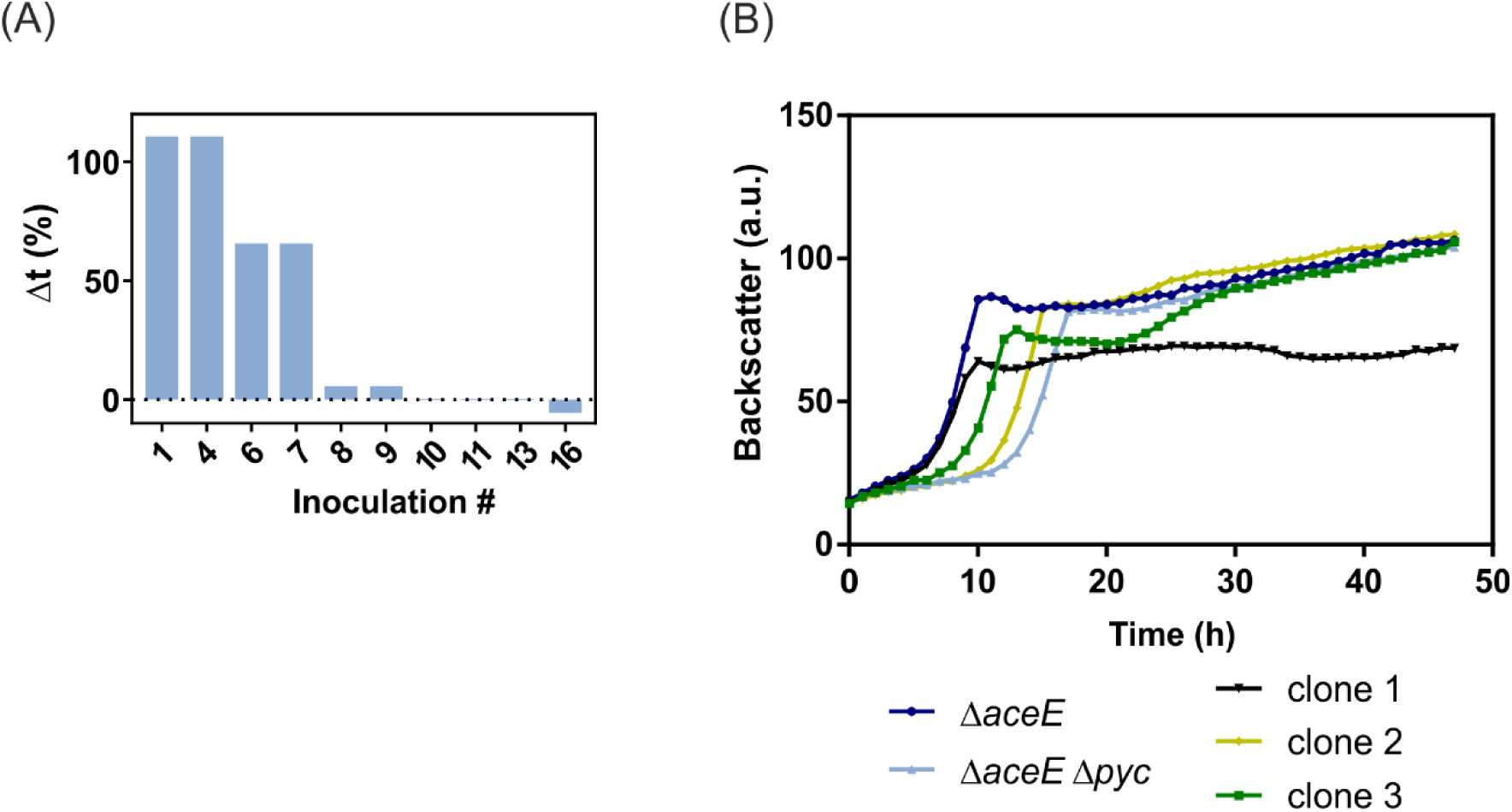
Adaptive laboratory evolution of *C. glutamicum* Δ*aceE* Δ*pyc*. Growth analysis of single inoculation steps obtained from the adaptive laboratory evolution (ALE) approach, which was performed with the strain Δ*aceE* Δ*pyc* in the presence of glucose and acetate. Growth curves are shown based on the backscatter measurements in a microtiter cultivation system. Symbols represent the backscatter means. (A) For growth analysis on the population level, glycerol stocks which were prepared during the ALE experiment, were used for the inoculation of a first pre-culture in BHI supplemented with 51 mM acetate. The second pre-culture in CGXII containing 154 mM acetate was then used for inoculation of the main culture in CGXII containing 154 mM acetate and 222 mM glucose (start OD_600_ of 1). Shown is the development of the relative lag phase duration after repetitive inoculations of Δ*aceE* Δ*pyc* in media containing glucose and acetate. The difference between first doubling time of Δ*aceE* and Δ*aceE* Δ*pyc* was set to 100%. For each inoculation, the difference of the lag phase compared to Δ*aceE* was calculated and is given in percent. (B) Single clones were isolated from batch number 7 (clone 1, black), 8 (clone 2, yellow) and 10 (clone 3, green) and were cultivated as mentioned above. Δ*aceE* (dark blue) and Δ*aceE* Δ*pyc* (light blue) served as control. These single clones were further analyzed by genome sequencing (Table 7).

**Table 7:**
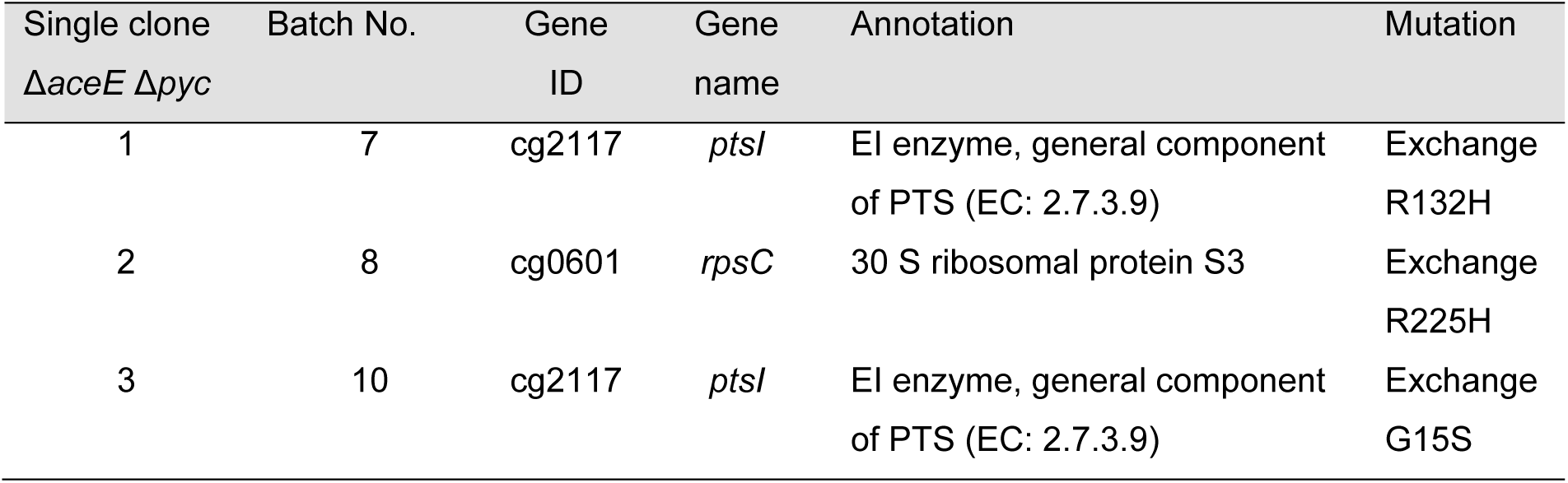
Key mutations identified in adaptive laboratory evolution experiment of *C. glutamicum* Δ*aceE* Δ*pyc*.

In summary, it was indeed possible to eliminate the observed lag phase of strain Δ*aceE* Δ*pyc* ascribed to glucose sensitivity by an adaptive laboratory evolution approach. Especially, the mutations identified in the *ptsI* gene are in agreement with other experiments emphasizing the high selective pressure exerted by glucose on PCx-deficient strains when the intracellular HCO_3_^-^ levels are limiting.

## Discussion

During growth on glucose, the pyruvate dehydrogenase complex catalyzes a central metabolic reaction contributing to the intracellular CO_2_/HCO_3_^-^ pool. In this study, we systematically perturbed the intracellular CO_2_/HCO_3_^-^ pool of PDHC-deficient *C. glutamicum* strains to monitor the impact on growth and anaplerotic flux. Even the PDHC-deficient strain *C. glutamicum* Δ*aceE* features a slightly elongated lag phase on glucose-acetate mixtures compared to the wild type, while no difference in growth is observed when strains are growing on gluconeogenetic substrates, like acetate. This is remarkable since this strain catabolizes only a minor fraction of the provided glucose in the exponential growth phase, but rather turns to the production of L-valine and L-alanine from glucose in the stationary phase (11, 12). The growth defect of *C. glutamicum* Δ*aceE* was, however, successfully complemented by the addition of HCO_3_^-^ which had no significant effect on growth of wild type cells, therefore, hinting towards problems caused by an impaired anaplerotic flux in this strain background. By comparing the CO_2_ production rates of the *C. glutamicum* Δ*aceE* strain and the wild type, Bartek et al. could show in a previous study, that strain Δ*aceE* excretes only around 0.83 mmol g^-1^ h^-1^ of CO_2_, while the wild type excretes ca. 1.65 mmol g^-1^ h^-1^ CO_2_ (30). The lower CO_2_ production rates of the PDHC-deficient strain already indicate a significant impact on the intracellular CO_2_/HCO_3_^-^ pool affecting metabolic flux distribution especially at low cell densities when CO_2_/HCO_3_^-^ is limiting.

The anaplerotic node comprises the essential link between glycolysis/gluconeogenesis and the TCA cycle (21). In contrast to most other organisms, *C. glutamicum* possesses both anaplerotic carboxylases, PEPCx (encoded by *ppc*) and PCx (encoded by *pyc*) catalyzing the anaplerotic bicarbonate-dependent reactions to yield oxaloacetate from PEP or pyruvate, respectively. In *C. glutamicum*, it was shown that these two enzymes can replace each other to a certain extent depending on the intracellular concentrations of the respective effectors for each enzyme. However, during growth on glucose, the biotin-containing PCx contributes to the main anaplerotic activity of 90% compared to PEPCx (24, 33). In our study, deletion of the *pyc* gene resulted in a severely elongated lag phase (>15 hours) of Δ*aceE* Δ*pyc* strains when glucose was present in the medium. Retarded growth of Δ*pyc* strains have been reported previously and, in line with our data, Blombach and co-workers also reported a severe growth defect of this strain under low CO_2_/HCO_3_^-^ levels (24, 29). This effect appeared to be even more severe when the cells were grown in microtiter plates under CO_2_-stripping conditions (low culture volume, high air mixing) compared to growth in higher culture volumes (Fig. S6). The PDHC-deficient mutant strain *C. glutamicum* Δ*aceE* Δ*pyc* was found to react very sensitively towards small changes in the intracellular bicarbonate availability and was therefore used as a test platform to systematically assess the impact of perturbations affecting the intracellular CO_2_/HCO_3_^-^ pool. While the addition of HCO_3_^-^, the increase of the pH as well as higher culture volumes rescued the strain, mutation of the urease accessory protein (*ureD-*E188*), which lowers the intracellular CO_2_/HCO_3_^-^ pool, resulted in an even more severe phenotype. It was not possible to restore growth if the second anaplerotic gene *ppc* was deleted as well, confirming that the abovementioned measures fostered the flux over the remaining anaplerotic reaction catalyzed by PEPCx.

The mutation in *ureD* was revealed by a previous biosensor-driven evolution approach selecting for mutations enhancing L-valine production in *C. glutamicum* (34). Here, inactivation of the urease enzyme by the mutation *ureD-*E188* reduced the anaplerotic flux via PCx resulting in an increased precursor availability of pyruvate-derived products like L-valine and L-alanine (34). In the present study, we further confirmed this finding by complementation with HCO_3_^-^ counteracting the effect of the *ureD-*E188* mutation. In another study by Blombach et al., lower CO_2_/HCO_3_^-^ levels also triggered enhanced production of L-valine and L-alanine of *C. glutamicum* during shake flask cultivation (29). However, a positive impact of anaplerotic reactions has been confirmed for other, for example, oxaloacetate-derived products like L-lysine. Here, attempts to overexpress *pyc* or the introduction of deregulated variants significantly improved L-lysine production (42, 43). However, in spite of great efforts and successes in the development of L-lysine strains, the impact of altered CO_2_/HCO_3_^-^ levels has not been systematically assessed so far.

Residual glucose consumption of PDHC-deficient *C. glutamicum* strains has already been reported in previous studies (11, 12, 30). The results of this work, however, emphasize that mutants lacking the major anaplerotic route via PCx are under strong evolutionary pressure in the presence of glucose. Although strains would be capable to grow on the acetate supplied in the growth medium, increasing levels of glucose resulted in a severely elongated lag phase (up to >40 hours with 250 mM) of *C. glutamicum* Δ*aceE* Δ*pyc.* Besides the abovementioned efforts to increase the CO_2_/HCO_3_^-^ pool, also the deletion of the *ptsG* gene itself effectively restored growth on glucose, but not on other PTS substrates. This indicates that this strain is sensitive to glucose, but does not have issues with regulatory functions of PtsG known from e.g. *E. coli* (44, 45).

Furthermore, among the three key mutations identified in the ALE experiment, two SNPs obtained from independent cell lines were located within the *ptsI* gene, encoding the EI component of the PTS system. These findings clearly highlight the problems of residual glucose consumption in strains featuring an impaired anaplerotic flux. One reason for the observed growth phenotype might be stress resulting from the intracellular accumulation of sugar phosphates. In *Escherichia coli*, sugar phosphate stress can be caused by accumulation of any sugar phosphates due to a block in glycolysis (46), e.g. by mutations in *pgi* or *pfkA* (47), or due to excessive glucose uptake caused e.g. by overexpression of *uhpT* encoding a sugar phosphate permease (48). The resulting metabolic imbalance causes growth inhibition. For example, in *C. glutamicum*, accumulation of several sugar phosphates, like fructose-1,6-bisphosphate or PEP, inhibits the isocitrate lyase (35). To counteract this stress, *E. coli* triggers SgrR which consequently activates SgrS by an unknown signal. The transcription factor SgrS then prevents further uptake of glucose by downregulation of *ptsG* (46, 49, 50). This is in line with the finding of our study, showing a slight downregulation of *ptsG* upon resumed growth of strain Δ*aceE* Δ*pyc* (Table 3). This reduction of glucose uptake via the PTS system which converts PEP to pyruvate during glucose transport also contributes to an increase in PEP availability for the remaining anaplerotic reaction catalyzed by PEPCx. Further, this is in line with an upregulation of *iolT1*, which ensures minor usage of glucose without PEP depletion (40, 41). In this context, also the downregulation of the *pyk* gene in strain Δ*aceE* Δ*pyc* can be interpreted as a potential adaptation to increase the PEP pool fostering anaplerotic flux.

During growth on gluconeogenic carbon sources - like acetate - the transcriptional repressor SugR represses the expression of the PTS genes in *C. glutamicum*, including the glucose-specific *ptsG* gene, but also *ptsF*, *ptsS* and general components as *ptsI* and *ptsH* (51–53). Derepression appears to be triggered by the accumulation of fructose-6-phosphate (F6P), which is generated from glucose 6-phosphate entering glycolysis. Consequently, F6P accumulation leads to an increase of glucose consumption rates resulting in parallel catabolization of glucose and acetate (15). The fact that the PDHC-deficient strains are also subject to regulation of SugR was demonstrated by studies showing that Δ*aceE* Δ*sugR* strain leads to up to 4-fold higher glucose consumption rates (54).

In aerobic glucose-based bioprocesses the endogenous production of CO_2_ is typically sufficiently high to promote microbial growth even at low cell densities. However, it is a well-known fact that anaerobic growth of some bacteria, like *E. coli*, requires the external supply of CO_2_/HCO_3_^-^ to avoid long lag phases (26, 55, 56). The critical impact of CO_2_/HCO_3_^-^ levels may become especially evident under conditions when the endogenous CO_2_ production rate becomes limiting. This was nicely demonstrated by a recent study of Bracher et al. who could show that long lag phases of engineered, but non-evolved *Saccaromyces cerevisiae* strains during xylose fermentation could be avoided by sparging the bioreactor cultures with CO_2_/N_2_ mixtures (57). Alternatively, the addition of L-aspartate, whose transamination provides oxaloacetate refueling the TCA, completely abolished the long adaptation phase of the respective yeast strains. In line with these findings, a recent ^13^C-flux analysis with *E. coli* revealed that a considerable turnover of lipids via β-oxidation appears to be required for growth on xylose to enhance the intracellular CO_2_/HCO_3_^-^ pool to a growth-promoting level enabling anaplerotic flux (58). These findings are in nice agreement with our study showing that the elongated lag phase of strain *C. glutamicum* Δ*aceE* Δ*pyc* can be eliminated by different measures enhancing either the intracellular CO_2_/HCO_3_^-^ level or by refueling of the TCA cycle with different intermediates, like citrate, malate or succinate. Overall, oxaloacetate depletion appeared to represent a key issue causing the delayed growth of the Δ*aceE* Δ*pyc* strain. Our data revealed that this is caused by an impaired anaplerotic flux on glucose due to low intracellular CO_2_/HCO_3_^-^ levels as well as reduced activities of glyoxylate enzymes.

Altogether, these results emphasize the important impact of the intracellular CO_2_/HCO_3_^-^ pool for metabolic flux distribution, which is especially relevant in engineered strains suffering from lower endogenous CO_2_ production rates as exemplified by PDHC-deficient strains in this study, but also by the performance of pentose-fermenting yeast and *E. coli* strains (57, 58). Consequently, the lack of an important by-product, like CO_2_ released by the PDHC complex, may have a significant impact on cellular metabolism and growth, especially on glycolytic substrates demanding a high flux via the anaplerotic reactions.

## Material and methods

### Bacterial strains and growth conditions

All bacterial strains and plasmids used in this study are listed in Table 1. Mutant strains are based on the *Corynebacterium glutamicum* ATCC 13032 wild type strain (59).

Standard cultivation of *C. glutamicum* Δ*aceE* cells and derivatives were performed on BHI (brain heart infusion, Difco BHI, BD, Heidelberg, Germany) agar plates containing 51 mM potassium acetate (hereinafter named acetate) (ChemSolute, Th. Geyer, Stuttgart, Germany) at 30°C for two days. One single colony was picked and incubated for 8 to 10 h at 30°C in either 4.5 ml or 1 ml BHI containing 154mM acetate in reaction tubes or Deepwell-plates (VWR International, Pennsylvania, USA), respectively. First pre-cultures were used to inoculate second pre-cultures in CGXII minimal medium (60) supplemented with 154 mM acetate either as 10 ml culture in shake flasks or in 1 ml volume in Deepwell-plates. After overnight growth, a main culture was inoculated at an OD_600_ of 1 in CGXII media containing 154 mM acetate and either 222 mM D(+)-glucose monohydrate (Riedel-de Haën, Seelze, Germany) (further referred to as glucose), or any other carbon source as stated, e.g. D(-)-fructose (Sigma-Aldrich, Taufkirchen, Germany) (referred to as fructose), D(+)-sucrose (Roth, Karlsruhe, Germany) (referred to as sucrose), D-gluconic acid sodium salt (Sigma-Aldrich, Taufkirchen, Germany) (referred to as gluconate), D(-)ribose (Sigma-Aldrich, Taufkirchen, Germany) (referred to as ribose), citric acid monohydrate (Merck Millipore, Darmstadt, Germany) (referred to as citrate), H-Glu-Ala-OH (Bachem AG, Bubendorf, Germany) (referred to as Glu-Ala), succinic acid (Sigma-Aldrich, Taufkirchen, Germany) (referred to as succinate) or L-malic acid (Merck Millipore, Darmstadt, Germany) (referred to as malate). For elevating the extracellular availability of bicarbonate, 100 mM potassium HCO_3_^-^ (Merck Millipore, Darmstadt, Germany) was added to the CGXII-basis solution, which was sterile-filtered before further use. In order to analyze the effect of different pH levels, either HCl for lowering or KOH for increasing the pH was added to the CGXII-basis, which was sterile-filtered afterwards. For cultivations in the presence of TCA cycle-filling/refueling substrates, 50 mM citric acid monohydrate (citrate), 3 mM H-Glu-Ala-OH dipeptide (Glu-Ala), 100 mM succinic acid (succinate) or 100 mM L-malic acid (malate) was used. In experiments where gluconate or ribose were used as carbon source, 100 mM D-gluconic acid sodium salt or 100 mM D-ribose were added, respectively. Biomass formation was monitored during cultivation in shake flasks by measuring OD_600_ or by measuring backscatter values during microtiter plate cultivation (section 2.2). Where necessary, 25 µg/ml kanamycin was also supplemented.

For the purpose of cloning and plasmid isolations, *Escherichia coli* DH5α strains were grown in shake flasks in Lysogeny broth (LB) medium at 37°C directly inoculated from a glycerol stock or from an LB-agar plate in shake flasks at 37°C. If necessary for selection, 50 µg/ml kanamycin was also supplemented.

### Microtiter plate cultivation

Online monitoring of growth and/or pH was performed in 48-well microtiter Flower Plates® (m2p-labs GmbH, Baesweiler, Germany) sealed with sterile breathable rayon films (VWR International, Pennsylvania, US) in a BioLector® microtiter cultivation system (m2p-labs GmbH, Baesweiler, Germany) (61). The cultivation conditions were adjusted as described earlier (62) and biomass formation was recorded each 15 min as the backscattered light intensity (light wavelength 620 nm; signal gain factor of 20) for 24 to 72 h at 30°C and 1,200 rpm. pH measurements were performed with 48-well microtiter Flower Plates® being equipped with pH optodes. Obtained data was evaluated using the software BioLection (m2p-labs, Baesweiler, Germany) and Graph Pad Prism (Graphpad Prism 7 Software, Inc., California, USA).

### Adaptive laboratory evolution

The adaptive laboratory evolution (ALE) of *C. glutamicum* Δ*aceE* and Δ*aceE* Δ*pyc* was performed in Deepwell-plates (VWR International, Pennsylvania, US) in a main culture of 1 ml CGXII media containing 154 mM acetate and 222 mM glucose or solely 154 mM acetate (as control without selection pressure) adjusted to an OD_600_ of 1. Cells were cultivated for 2-3 days before the next generation was inoculated starting at an OD_600_ of 1 and cultivated again for 2-3 days. After each generation step, glycerol stocks of cultures were prepared (20% glycerol) and stored at −80°C allowing growth analysis and DNA sequencing of individual inoculations. In total, 16 serial transfers were analyzed.

### Cloning techniques and recombinant DNA work

Standard molecular biology methods were performed according to J. Sambrook and D. W. Russell (63). *C. glutamicum* ATCC 13032 chromosomal DNA was used as template for PCR amplification of DNA fragments and prepared as described earlier (64). DNA fragment and plasmid sequencing as well as synthesis of oligonucleotides was performed by Eurofins Genomics (Ebersberg, Germany).

For the construction of plasmids (Table S2), DNA fragments were amplified using the respective oligonucleotides (Table S1) and enzymatically assembled into a vector backbone according to (65).

To achieve genomic deletion of *pyc, ptsG, ppc* and *pgi*, two-step homologous recombination using the pK19*mobsacB*-system (66) was implemented. The suicide plasmids (compare Table 2 and Table S2) were isolated from *E. coli* cells using the QIAprep spin miniprep kit (Qiagen, Hilden, Germany). Electrocompetent *C. glutamicum* Δ*aceE* and Δ*aceE* Δ*pyc* cells were transformed with these plasmids by electroporation (67). The first and second recombination events were performed and verified as described in previous studies (68). The deletion of *pyc, ptsG, ppc* and *pgi* was reviewed by amplification and sequencing using primers shown in Table S1.

### Measurement of glucose concentrations

To measure the glucose concentration of the culture media at different time points, cultivation was performed in 50 ml in shaking flasks. During cultivation, 0.5 ml samples were taken every 3 h and centrifuged (16,000 x g). Supernatant was collected and stored at −20°C until use.

The measurement of the actual glucose concentration was performed using D-Glucose UV-Test Kit (r-biopharm, Darmstadt, Germany). Performance and calculations according to manufacturer’s instructions were done, while absorption was measured at 340 nm.

Further calculations concerning the glucose uptake rates could be done based on the obtained glucose concentrations and OD_600_ values. According to (18), Formula 1 was used:

**Formula 1:** Determination of glucose concentration in nmol gDW^-1^ h^-1^

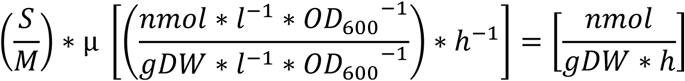

While S represents the slope of the glucose concentration versus OD_600_ [nmol * l^-1^ * OD_600_^-1^], M is the correlation between dry weight and OD_600_ [g dry weight * l^-1^ * OD_600_^-1^], and µ is the growth rate [h^-1^]. According to A. Kabus et al. (69), an OD_600_ of 1 corresponds to 0.25 g dry weight l^-1^, so that this value was used as M throughout this calculations.

### Quantification of amino acid production

Using ultra-high performance liquid chromatography (uHPLC), amino acids were quantified as *ortho*-phthaldialdehyde derivatives by automatic pre-column derivatization. Separation of derivatives by reverse-phase chromatography was performed on an Agilent 1290 Infinity LC ChemStation (Agilent, Santa Clara, USA) equipped with a fluorescence detector. As eluent for the Zorbax Eclipse AAA 3.5 micron 4.6 x 7.5 mm column (Agilent, Santa Clara, USA), a gradient of Na-borate buffer (10 mM Na_2_HPO_4_; 10 mM Na_2_B_4_O_7_, pH 8.2; adapted to operater’s guide) and methanol was applied. Prior to analysis, samples were centrifuged for 10 min at 16,000 x g and 4°C and diluted 1:100.

### Monitoring of cellular proliferation by cell staining

For staining of proliferating cells, the PKH67 green fluorescent cell linker kit for general cell membrane labeling (Sigma Aldrich, Munich, Germany) was used and the protocol was adapted to O. Kotte et al. (36). From an exponentially growing pre-culture in CGXII minimal medium containing 222 mM glucose and 154 mM acetate, 1.5 x 10^9^ cells were harvested by centrifugation for four minutes at 4000 *g* and 4°C. Then, the cells were washed again in 5 ml ice-cold CGXII basic solution, without MgSO_4_, CaCl_2_, biotin, trace elements and protocatechuic acid. For staining, the cell pellet was resuspended in 500 µl dilution buffer C (Sigma Aldrich, Munich, Germany) at room temperature and a freshly prepared mixture of 10 µl PKH67 dye (Sigma Aldrich, Munich, Germany) and 500 µl dilution buffer C was added. Subsequently, cells were incubated for three minutes at room temperature and afterwards, 4 ml ice-cold filtered CGXII basic solution containing 1% (w/v) bovine serum albumin, 1 mM MgSO_4_ and 0.1 mM CaCl_2_ were added. Then, the cells were centrifuged for four minutes at 4000 *g* and 4°C and the cell pellet was washed twice. Finally, the cells were resuspended in CGXII minimal medium containing 222 mM glucose and 154 mM acetate and cultivated in a microtiter cultivation system.

### Flow cytometry

Flow cytometric (FC) analyzes were conducted on a FACSAria II flow cytometer (Becton Dickinson, San Jose, USA) equipped with a blue solid state laser (488 nm excitation). Forward-scatter characteristics (FSC) and side-scatter characteristics (SSC) were recorded as small-angle and orthogonal scatters of the 488-nm laser, respectively. PKH67 fluorescence was detected using a 502-nm long-pass and a 530/30-nm band-pass filter set. FACS-Diva software 6.0 was used to monitor the measurements. During analyzes, thresholding on FSC was applied to remove background noise. For FC analyzes, PKH67-stained culture samples were diluted to an OD_600_ of 0.05 in FACSFlow™ sheath fluid buffer (BD, Heidelberg, Germany). The analysis software FlowJo V.10.0.8 was used to visualize and evaluate the data (Tree Star, Ashland, USA).

### DNA microarrays

For analysis of the transcriptome, *C. glutamicum* Δ*aceE* and Δ*aceE* Δ*pyc* were cultivated in triplicates as described in 2.1 in 50 ml CGXII containing 154 mM acetate and 222 mM glucose in shake flasks. After reaching exponential phase at an OD_600_ of around 12-15, the cell suspension was harvested by centrifugation (4256 *g*, 10 min, 4°C). The resulting pellet was directly frozen in liquid nitrogen and stored at −80°C. RNA preparation, cDNA synthesis as well as microchip hybridization, scanning and evaluation were performed as described by previous studies (70).

### GC-ToF-MS analysis

For analysis of the metabolome, samples of *C. glutamicum* Δ*aceE* in exponential phase as well as Δ*aceE* Δ*pyc* in stationary and exponential phase were taken and cell disruption was accomplished using hot methanol. Further sample preparation, derivatization, MS operation and peak identification was accomplished according to N. Paczia et al. (71) in an Agilent 6890N gas chromatograph (Agilent, Santa Clara, USA) coupled to a Waters Micromass GCT Premier high resolution time-of-flight mass spectrometer (Waters, Milford, USA). Identification of known metabolites was accomplished using in-house database JuPoD, the commercial database NIST17 (National Institute of Standards and Technology, US) and the database GMD (MPI of Molecular Plant Physiology, Golm) (72).

### Whole-genome sequencing

In order to sequence the whole genome of *C. glutamicum* Δ*aceE* Δ*pyc* mutants from the ALE experiment using next-generation sequencing (NGS), genomic DNA was prepared using the NucleoSpin® Microbial DNA Kit (Macherey-Nagel, Düren, Germany) according to manufacturer’s instructions. Consequently, the concentrations of the purified genomic DNA were measured using Qubit Fluorometer 2.0 (Invitrogen, California, USA) according to manufacturer’s instructions. Overall, 4 µg purified genomic DNA was employed for the preparation of genome sequencing using TruSeq® DNA Library Prep Kit and MiSeq Reagent Kit v1 (Illumia, San Diego, USA) according to manufacturer’s instructions. The sequencing run was performed in a MiSeq system (Illumina, San Diego, USA). Data analysis and base calling was performed with the Illumina instrument software. Consequent fastq output files were examined using CLC Genomics Workbench 9 (Qiagen Aarhus, Denmark).

## Acknowledgements

We thank Jochem Gätgens for performing GC-ToF analyses, as well as Bastian Blombach and Stephan Noack for fruitful discussions. The authors acknowledge financial support by the Helmholtz Association (grant W2/W3-096).

